# Evolution of transposable element-derived enhancer activity

**DOI:** 10.1101/2022.03.16.483999

**Authors:** Alan Y. Du, Xiaoyu Zhuo, Vasavi Sundaram, Nicholas O. Jensen, Hemangi G. Chaudhari, Nancy L. Saccone, Barak A. Cohen, Ting Wang

## Abstract

Many transposable elements (TEs) contain transcription factor binding sites and are implicated as potential regulatory elements. However, TEs are rarely functionally tested for regulatory activity, which in turn limits our understanding of how TE regulatory activity has evolved. We systematically tested the human LTR18A subfamily for regulatory activity using massively parallel reporter assay (MPRA) and found AP-1 and C/EBP-related binding motifs as drivers of enhancer activity. Functional analysis of evolutionarily reconstructed ancestral sequences revealed that LTR18A elements have generally lost regulatory activity over time through sequence changes, with the largest effects occurring due to mutations in the AP-1 and C/EBP motifs. We observed that the two motifs are conserved at higher rates than expected based on neutral evolution. Finally, we identified LTR18A elements as potential enhancers in the human genome, primarily in epithelial cells. Together, our results provide a model for the origin, evolution, and co-option of TE-derived regulatory elements.

## Introduction

Changes in gene regulation have long been implicated as crucial drivers in evolution^1^. Since the discovery of the SV40 enhancer element, enhancers have emerged as one of the major classes of cis-regulatory sequences that can modulate gene expression^2, 3^. Due to several unique properties, enhancers have emerged as excellent candidates upon which evolution can act. Enhancers are often active depending on cellular context like cell type or response to stimuli. This modularity can minimize functional trade-offs and allows selection to act more efficiently^4^. Furthermore, redundant enhancers, or “shadow” enhancers, provide robustness in gene regulatory networks and may allow for greater freedom to develop new functions^5, 6^.

The development of massively parallel reporter assays (MPRAs) has greatly accelerated our understanding of enhancers by facilitating simultaneous testing of thousands of DNA sequences^7–10^. MPRAs have been used to probe the enhancer potential of sequences underlying various epigenetic marks^11^, dissect enhancer logic through tiling and mutagenesis^9, 12, 13^, and decipher the effects of naturally occurring sequence variants^8, 14–16^. Several studies have also employed MPRA to understand the evolution of fly and primate enhancers, revealing widespread enhancer turnover^17, 18^.

Transposable elements (TEs) are repetitive DNA elements that represent a rich source of genetic material for regulatory innovation^19^. In mammalian genomes, TEs have made substantial contributions to the collection of transcription factor binding sites^20–24^. These binding sites are often enriched within certain TE subfamilies, groups of similar TE sequences that are derived from a single ancestral origin. Individual copies of TE subfamilies can then be co-opted into gene regulatory networks such as in pregnancy and innate immunity^25, 26^. Overall, TEs make up a quarter of the regulatory epigenome in human^27^, and by some estimates, the majority of primate-specific regulatory sequences are derived from TEs^28, 29^. Despite these advances in the field, there remains a gap in knowledge of how TEs obtain regulatory activity and how this activity changes over the course of evolution.

As repetitive sequences, TEs offer a unique perspective into the evolution of cis-regulatory elements. One intrinsic limitation for evolutionary studies is that each enhancer has one ortholog per species barring duplication or deletion, which constrains the sample size for analysis. Within a TE subfamily, each TE is descended from a common ancestor, with each copy evolving mostly independently. This provides a large sample size to draw upon within even a single genome. To serve as a representative subfamily, we selected LTR18A which we previously identified to be enriched for MAFK transcription factor binding peaks and motifs^24^.

Here, we aim to investigate the evolution of regulatory potential in the LTR18A subfamily using MPRA. By using present day LTR18A sequences found across seven primate species, we computationally reconstruct ancestral sequences during LTR18A evolution across a span of roughly 75 million years. We apply tiling and motif-focused approaches to test reconstructed and present day LTR18A sequences for enhancer activity. Using natural sequence variations between LTR18A elements, we identify transcription factor binding sites that drive LTR18A enhancer activity and validate them through mutagenesis. By annotating enhancer activity for the root and intermediate ancestral LTR18A elements in our reconstructed phylogenetic tree, we investigate the origin of enhancer activity for the LTR18A family as well as key mutations that have led to changes in activity over time. Finally, we explore the influence of selection on LTR18A and the possibility of co-option in the human epigenome.

## Results

### Reconstruction of the LTR18A phylogenetic tree

In order to reconstruct the evolutionary history of the LTR18A subfamily, we first identified high confidence LTR18A elements in human and their orthologous elements in six other primate species. The LTR18A subfamily is found in the Simiiformes taxa^30^. From the Simiiformes, we obtained RepeatMasker annotations for human (hg19), chimpanzee (panTro4), gorilla (gorGor3), gibbon (nomLeu3), baboon (papAnu2), rhesus macaque (rheMac3), and marmoset (calJac3) genomes. Due to the similarity of the LTR18A, LTR18B, and LTR18C consensus sequences, we performed manual curation of hg19 LTR18A to select for LTR18A elements that are confidently assigned to the subfamily. Briefly, we filtered out LTR18A elements that could be aligned to either the LTR18B or LTR18C consensus, and we removed LTR18A elements that might be misannotated using paired LTRs (Methods). Following these criteria, 181 out of 198 LTR18A elements annotated by RepeatMasker are retained (Supplemental Table 1). Next, we found primate orthologs for each hg19 LTR18A element by using synteny^31^. LTR18A elements that correspond with multiple orthologs in the same genome, or vice versa, were excluded. Each hg19 LTR18A element with its primate orthologs were considered an ortholog set. We further selected for LTR18A pairs that have orthologs in chimpanzee, gorilla, and at least two of the four other primates. In the end, 46 (consisting of 23 pairs) LTR18A ortholog sets were chosen for ancestral reconstruction.

From our set of manually curated human LTR18A elements and their orthologs, we computationally reconstructed the LTR18A phylogenetic tree using a two-step process. Based on the unique characteristic of TEs to multiply by transposition and the presence of orthologous copies in different primate genomes, we split our reconstruction of LTR18A evolution into two phases corresponding to transposition and speciation (Figure 1A). For each of the 46 sets of LTR18A orthologs, we aligned orthologs using MAFFT and then reconstructed ortholog ancestor and intermediate sequences using PRANK^32, 33^. Then, using the ancestor sequences for the 46 LTR18A orthologs, we aligned and reconstructed the LTR18A subfamily ancestor as well as intermediates predating speciation. PRANK was chosen for ancestral sequence and phylogenetic tree reconstruction due to its ability to model insertions and deletions. However, PRANK tends to be biased towards insertions in our reconstruction. Thus, we manually curated sequences following PRANK reconstruction for both ortholog ancestors and subfamily ancestors (Methods).

**Figure 1:**
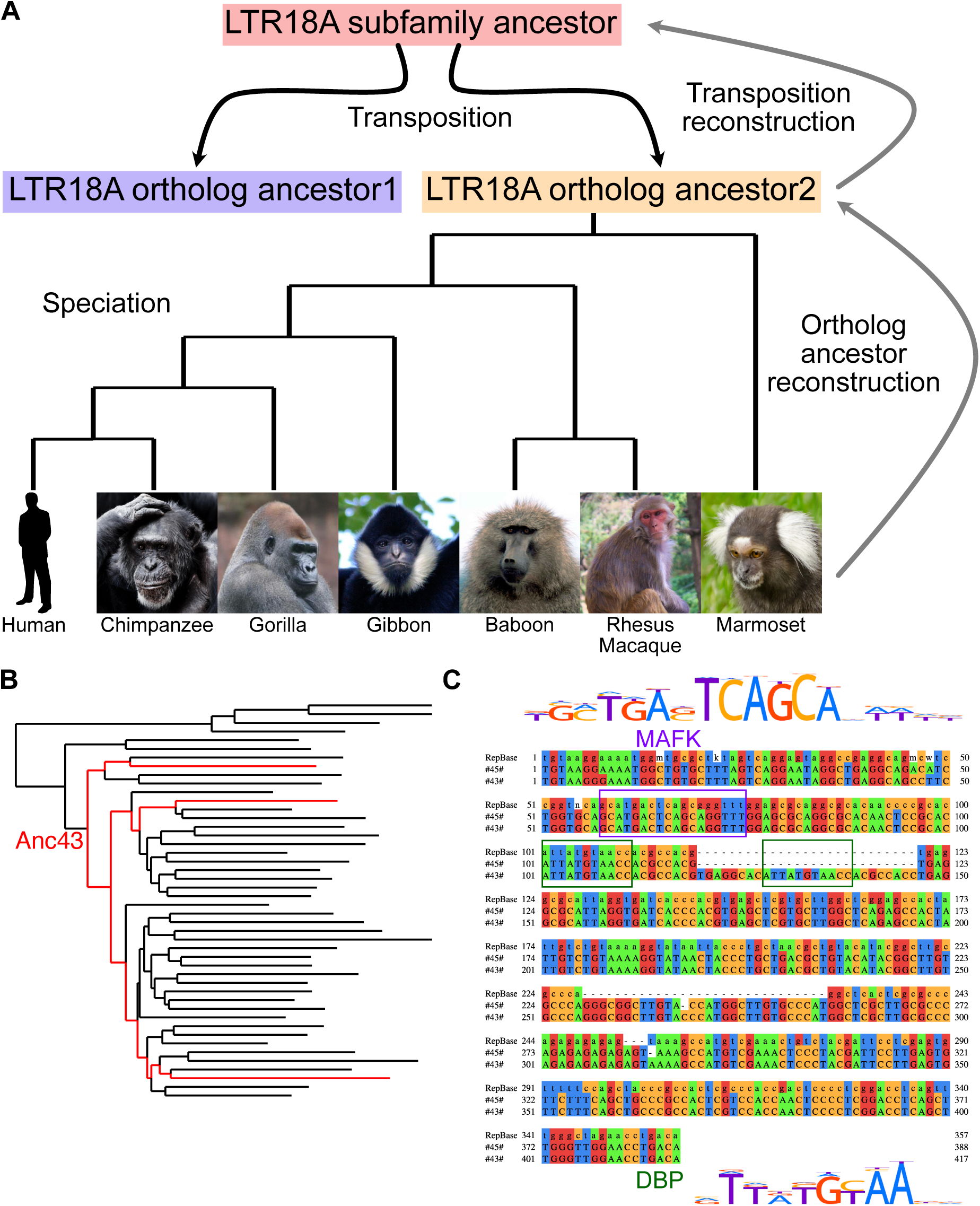
LTR18A ancestral reconstruction. A) Model of LTR18A evolution split into transposition and speciation phases. Computational reconstruction was performed for ortholog ancestors and transposition intermediates using PRANK. B) Phylogenetic tree for reconstructed transposition intermediates and ortholog ancestors at leaves. Ancestral node 43 (#43#) is labeled in red, as well as the edges to ortholog ancestors that contain the 27bp insert. C) Alignment of RepBase consensus, ancestral node 45 (#45#, subfamily ancestor), and ancestral node 43. Motifs in the sequences are boxed. DBP is shown to represent C/EBP-related motifs.

Next, we evaluated our reconstructed LTR18A sequences to see if they are consistent with those derived from other methods. TE consensus sequences are often used as a representation of the ancestral state of the subfamily. Excluding insertions and deletions, our reconstructed LTR18A subfamily ancestor has ∼5.9% substitution rate relative to the LTR18A consensus sequence, which is lower than the 16.1% subfamily average. This suggests that although we start from different elements and use different methodologies, both our reconstruction and the RepBase consensus are approaching each other. In addition to substitutions, our reconstructed ancestor also has ∼8.0% insertions compared to the consensus. The insertions appear to be caused by the consensus dropping bases if the majority of elements do not have the base in the alignment, as well as PRANK’s tendency to include insertions when alignable sequence is present in more than one element. The MAFK motif enriched in LTR18A was present in both our reconstructed subfamily ancestor and the RepBase consensus. Overall, the topology of our reconstructed phylogenetic tree resembles the tree generated from all hg19 LTR18A elements (Supplemental Figure 1). One feature of note occurs in node 43, two nodes from the root of the tree (Figure 1B). Relative to the subfamily ancestor, node 43 has a 27bp insertion that contains a C/EBP-related factor motif (Figure 1C). When we examined ortholog ancestor reconstructions for this insertion, three ortholog ancestors have an alignable 27bp insert, and the insertion is present in all present-day primate orthologs (Supplemental Figure 2). In hg19, 13/181 elements contain the insert. The insert-containing elements are spread throughout most of the hg19 LTR18A phylogenetic tree, which is consistent with a deep ancestral origin for the insert and occurrence in node 43 of our reconstruction. Additionally, the C/EBP motif is also found in the LTR18A consensus and enriched in the subfamily relative to genomic background. If the C/EBP motif is functionally important, the insertion of a second C/EBP motif could be an ancestral gain of function mutation. In conclusion, our reconstruction is able to generate a subfamily ancestor similar to the RepBase consensus and reveals evolutionary events that would otherwise be missed.

### Identification of important TFBS motifs in LTR18A enhancers

We designed our LTR18A MPRA library to assay elements at two resolutions (Figure 2). In one half, we synthesized motif-focused regions for 1225 LTR18A elements found across seven primate genomes, 280 ancestral reconstruction elements, and the RepBase consensus (Figure 2A). Specifically, we took the sequence of each element aligning to the first 160bp of our reconstructed ancestral node 43 (Methods). This allowed us to focus on the effects of sequence variation for both the MAFK motif and the C/EBP motif. In the other half of the library, we synthesized 160bp tiles at 10bp intervals of all pre-speciation ancestral reconstruction elements, ortholog ancestors and their present-day hg19 elements, and the LTR18A consensus (Figure 2B). We cloned LTR18A motif-focused regions and tiles upstream of a pGL4 vector with the hsp68 promoter (Figure 2C).

**Figure 2:**
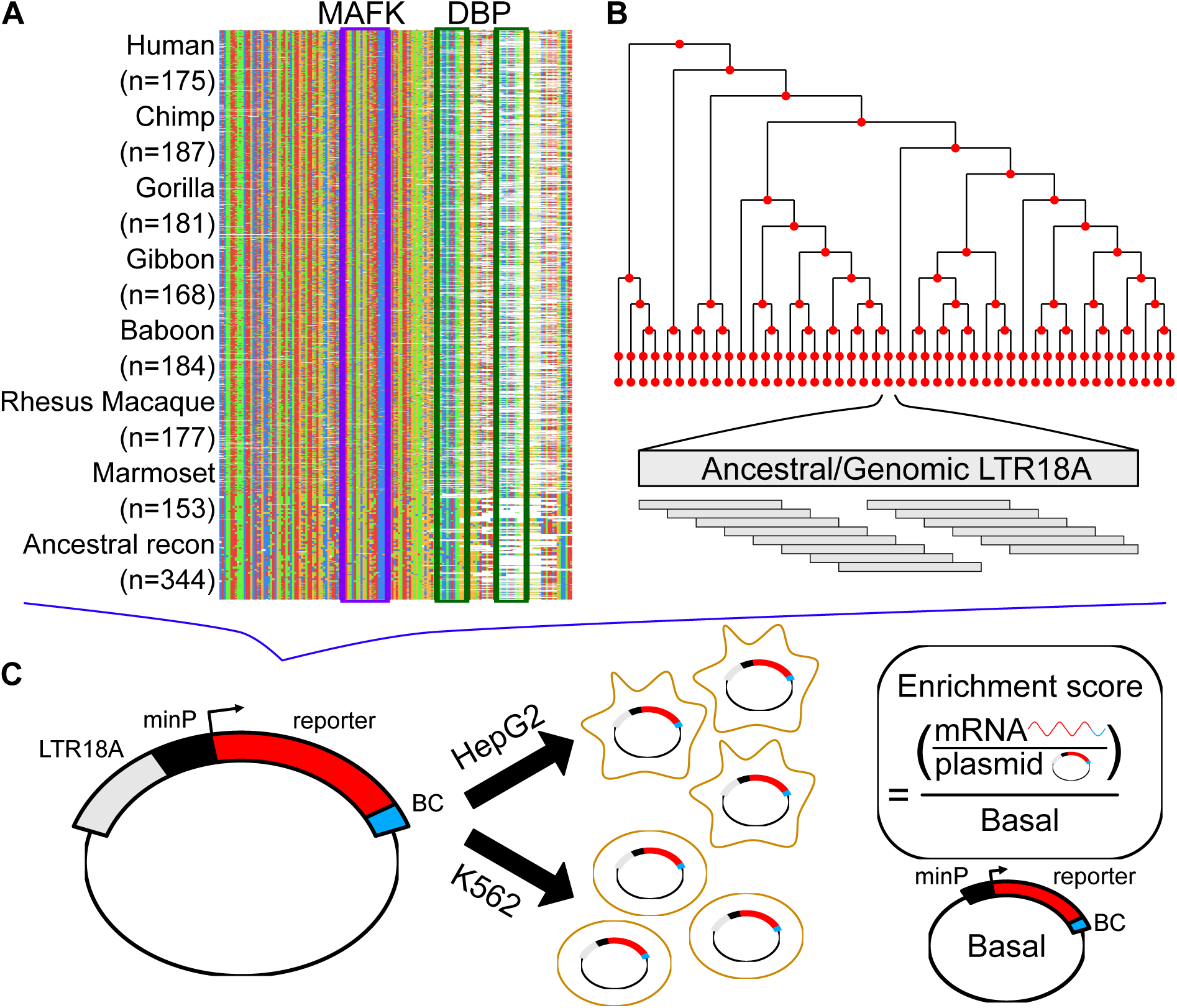
Schematic of MPRA. A) Sequence alignment of motif-focused regions to test primate and ancestral reconstructed LTR18A elements. MAFK and DBP motif regions are boxed. B) Tiling of ancestral and hg19 genomic LTR18A elements in reconstructed phylogenetic tree. All elements were tiled with 160bp tiles at 10bp intervals. C) Plasmid construct and enrichment score calculation. Each LTR18A fragment was integrated upstream of a minimal promoter (minP) and tagged with 10 unique barcodes (BC). The MPRA library was transfected into HepG2 and K562 cells. Enrichment scores are log_2_ ratios of RNA/DNA normalized to Basal.

To understand cell type effects, we tested LTR18A for enhancer activity in HepG2 and K562 cell lines. We calculated enrichment scores for each element by taking the log2 of the RNA over DNA ratio followed by normalization to the basal hsp68 promoter. Normalizing to the basal promoter allowed us to have the same reference point between cell lines. Active elements were defined as those with enrichment scores greater than 1, representing elements that increase transcription by greater than twofold. When we compare the distribution of enrichment scores for HepG2 and K562, we find that LTR18A elements are generally more active in HepG2 than K562 (Figure 3A). Out of 1506 motif-focused sequences tested, 1004 were classified as active in HepG2 while only 52 were classified as active in K562. For genomic LTR18A, 786 (123 from hg19) were active in HepG2 and 31 (4 from hg19) were active in K562. Enrichment scores are positively but poorly correlated between HepG2 and K562 despite high correlations between biological replicates (p<2.2e-16, Figure 3B, Supplemental Figure 3), implying differential sequence features required for enhancer activity between cell lines.

**Figure 3:**
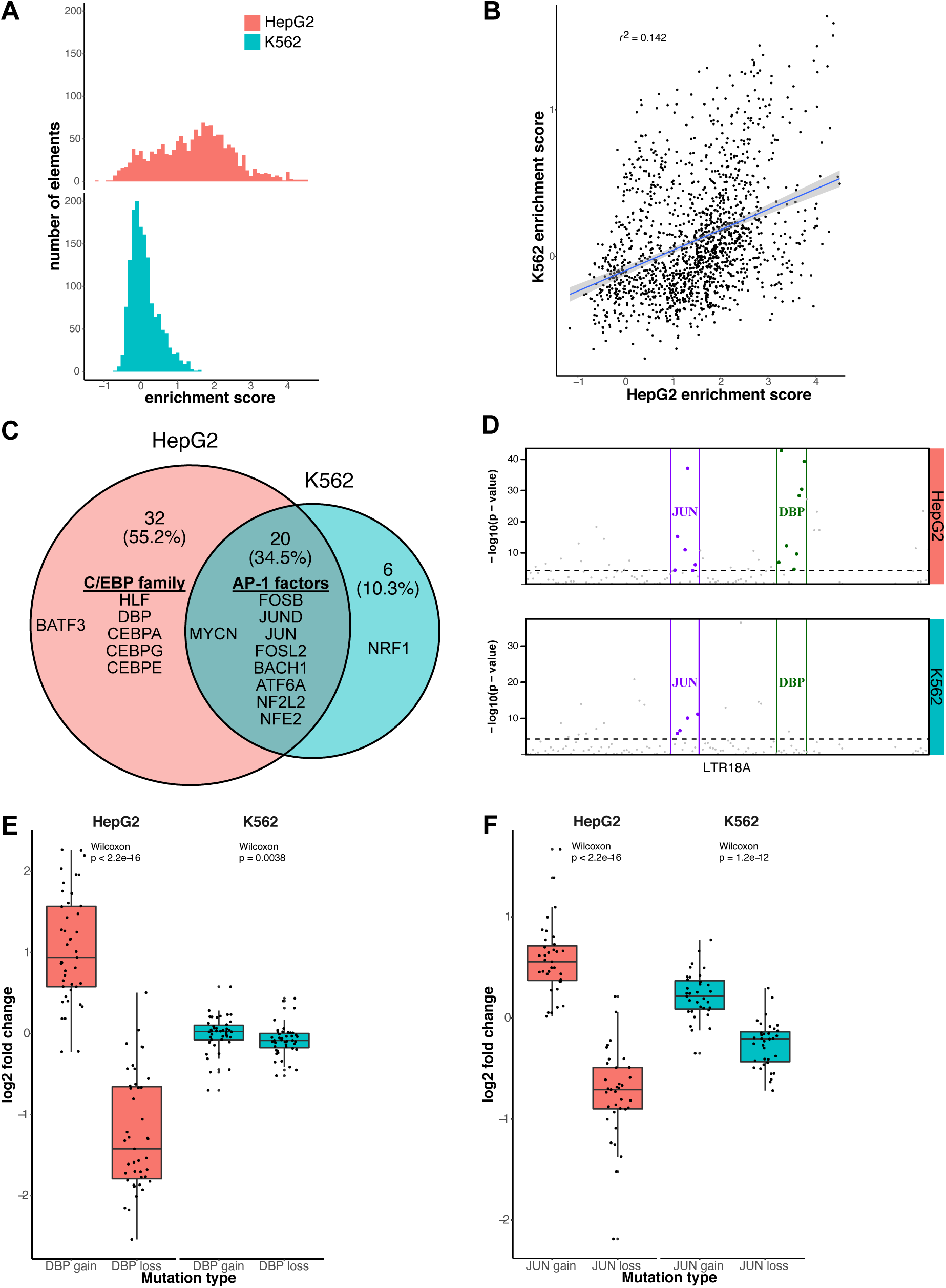
AP-1 motifs drive enhancer activity in HepG2 and K562 while C/EBP motifs are HepG2 specific. A) Distribution of enrichment scores of LTR18A motif focused regions in HepG2 and K562. B) Correlation of enrichment scores between HepG2 and K562. C) Overlap of motifs significantly associated with active LTR18A. Top 10 transcription factor motifs for both cell lines are displayed. AP-1 and C/EBP-related transcription factors are grouped. D) TEWAS significant nucleotides associated with active LTR18A. JUN and DBP motifs representing AP-1 and C/EBP-related motifs are boxed. Significant positions (p<5e-5, above dotted line) within the two motifs that are associated with active elements are highlighted. E) DBP mutagenesis effects on enhancer activity. F) JUN mutagenesis effects on enhancer activity. *P* values were derived from two-tailed Mann-Whitney *U* tests.

To identify important sequence features for enhancer activity, we took advantage of the natural sequence variation within LTR18A elements. Using AME motif enrichment analysis^34^, we asked if active elements were enriched for motifs compared to the rest of elements as background. Overall, 34.5% (20/58) motifs were enriched in active elements in both HepG2 and K562 (Figure 3C). Of the shared motifs, AP-1 (JUN, FOS, and ATF family) motifs were in the top 10 most enriched for both cell lines. Top 10 most enriched motifs that were cell line specific include the C/EBP family motifs and BATF3 for HepG2 and NRF1 in K562. As an orthologous method, we investigated if individual nucleotide positions are associated with enhancer activity. As this is analogous to genome-wide association studies (GWAS) but focused on sequence variation within a TE subfamily, which we term TE-WAS, we adapted the GWAS tool PLINK to find significant nucleotides^35, 36^. In HepG2, 6/11 JUN (AP-1) motif bases and 8/11 DBP (C/EBP family) motif bases are significantly associated with increased enhancer activity (Figure 3D). In K562, after we adjusted our cutoff for active elements to be an enrichment score of at least 0.5 to increase the number of active elements from 52 to 239, 4/11 JUN motif bases and 0/11 DBP motif bases are significantly associated with increased enhancer activity. In summary, both motif enrichment and TE-WAS approaches implicate AP-1 motifs as important to both HepG2 and K562 LTR18A enhancer activity while C/EBP-related motifs are HepG2-specific.

To validate the importance of C/EBP and AP-1 motifs to enhancer activity, we created targeted mutations in the motif regions of LTR18A elements. We chose DBP to represent the C/EBP family and JUN to represent the AP-1 family. We selected pairs of LTR18A orthologs of which one has the motif and the other does not by FIMO motif scanning^37^. For elements with the motif, we mutated the motif bases to low information nucleotides based on the PWM. For elements without the motif, we changed the motif aligned region to the consensus motif bases. To quantify the effect of motif mutations on enhancer activity, we took the log2 ratio of each motif mutated LTR18A sequence to its native sequence (Figure 3E, 3F). On average, DBP mutation gain and loss lead to a 2.07-fold increase and 2.36-fold decrease in enhancer activity respectively in HepG2. In contrast, the same DBP mutations have little effect in K562. JUN gain and loss lead to 1.49-fold increase and 1.68-fold decrease in HepG2 enhancer activity and 1.17-fold increase and 1.2-fold decrease in K562 enhancer activity. Both DBP and JUN mutagenesis results are consistent with our previous findings based on motif association.

### Evolution of LTR18A enhancer activity linked to sequence evolution

One of our primary goals was to understand how enhancer activity of LTR18A as a subfamily changed over time. To address this question, we synthesized 160bp tiles at 10bp intervals across each LTR18A ancestral sequence, ortholog ancestor, and hg19 element used in reconstruction (Figure 2B). After obtaining enrichment scores, we estimated nucleotide activity scores across each element to infer their relative effects on enhancer activity using the SHARPR software for MPRA tiling designs^12^. Due to overall low activity in K562, we focus on HepG2 for evolutionary analysis. When examining nucleotide activity scores across the length of our reconstructed LTR18A subfamily ancestor, we observe regions of increased activity over basal. The C/EBP and AP-1 motifs that we previously identified to be important for enhancer activity are embedded within the largest active region located near the start of the sequence (Supplemental Figure 6). Across LTR18A elements of our reconstructed phylogenetic tree, we were able to confirm that regions of increased SHARPR nucleotide activity were enriched for C/EBP and AP-1 motifs. As SHARPR nucleotide activity scores could discover the same biologically meaningful sequences as our previous analyses, we took the sum of activity scores across each LTR18A element and annotated them in our tree (Figure 4A). From a broad perspective, we were able to make several observations. First, the most divergent (leftmost) lineage on the tree loses enhancer activity early, and enhancer activity throughout the lineage remains low to the present day (Figure 4C). The low regulatory activity of the lineage could be linked to its relatively low rate of expansion (27/181 LTR18A elements in the lineage) (Supplemental Figure 7). This low activity lineage contrasts with the rest of the tree where evolutionary intermediates exhibit relatively high activity followed by less active elements at ortholog ancestor and present-day elements. Indeed, the overall trend appears to be that enhancer activity decreases over time, as shown by the decrease in mean SHARPR sum with increasing divergence from the LTR18A subfamily ancestor (Figure 4B). On the other hand, there is an increase in activity in the middle lineages, some of which persists to the ortholog ancestors and present-day elements (Figure 4D). Finally, enhancer activity of present day hg19 LTR18A elements and their corresponding ortholog ancestors are positively correlated with mostly small differences in activity, implying that post-speciation evolution has had small effects on regulatory potential overall (Supplementary Figure 8).

**Figure 4:**
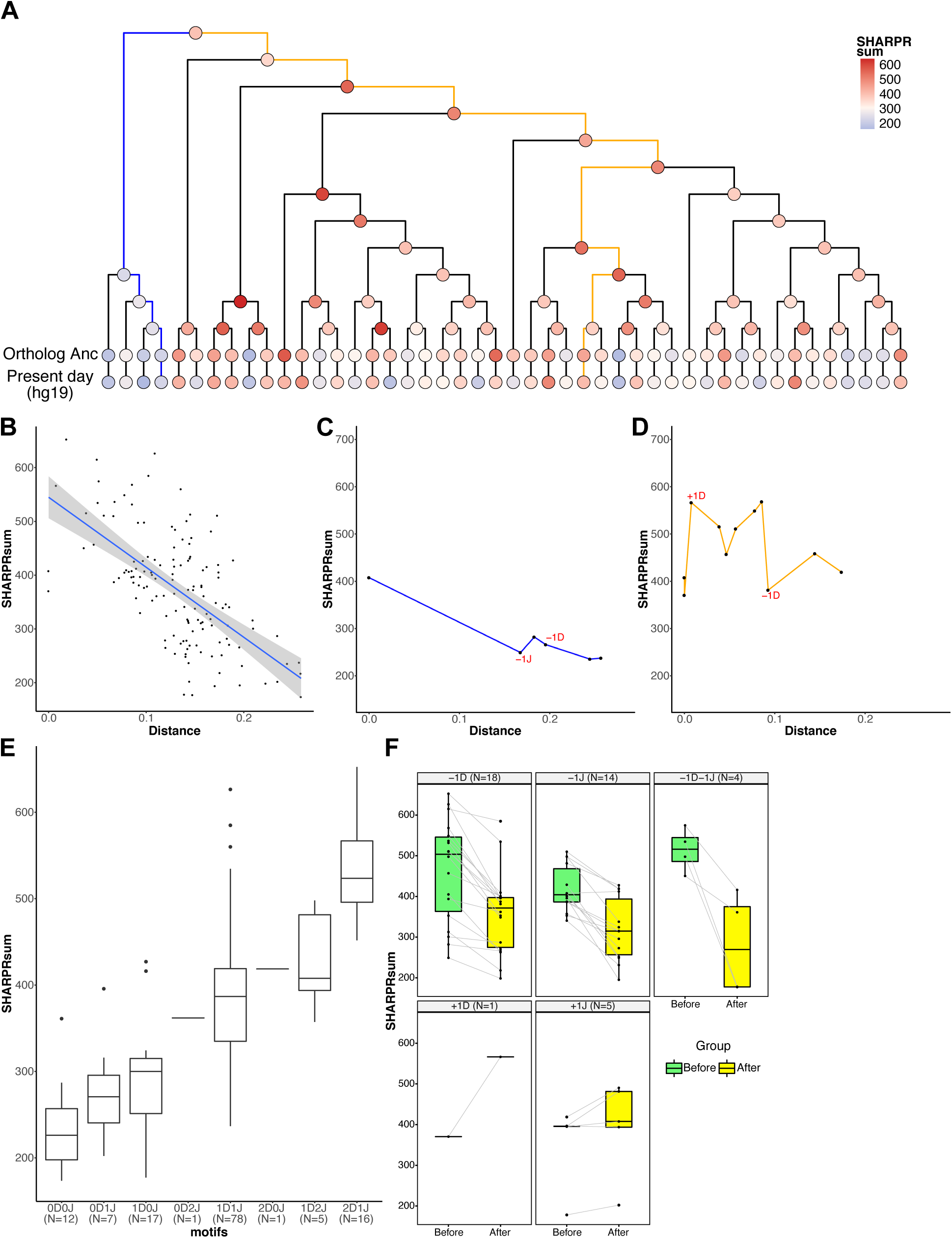
Evolution of regulatory activity in LTR18A in HepG2. A) Phylogenetic tree of reconstructed ancestral LTR18A annotated at each node/element with the sum of SHARPR nucleotide activity scores. B) Correlation of SHARPR sum and distance (substitution rate) from subfamily ancestor for each LTR18A in the phylogenetic tree. C) Example of regulatory activity evolution along the blue path in A. Motif changes are labeled in red (D = DBP, J = JUN). D) Same as C, but for the orange path in A. E) Distribution of SHARPR sums for phylogenetic tree elements separated by DBP and JUN motif content. F) Motif associated changes in SHARPR sum. Each motif change in the phylogenetic tree is shown with the before and after motif change SHARPR sums connected by a line.

To further investigate why enhancer activity changes in our LTR18A tree, we looked at differences in C/EBP and AP-1 motif presence using DBP and JUN as representatives. When elements are categorized by the number of DBP and JUN motifs, the number of motifs is positively correlated with SHARPR sum (Figure 4E). Furthermore, DBP or JUN loss correlates with a decrease in SHARPR sum, with rare motif gains generally corresponding to increased SHARPR sums (Figure 4F). Due to the significance of the DBP motif, we evaluated ancestral node 43 as the sole evolutionary intermediate that gained a second motif through an insertion event (Figure 1B). The motif gain leads to an increase in SHARPR sum of ∼39%, which is similar to the average effect size of the DBP motif (∼38%). This effect is validated by mutagenesis of our LTR18A subfamily ancestor and consensus to have the same 27bp insertion (34% and 32% increase respectively) as well as ablation of the second DBP motif in ancestral node 43 (41% decrease). In summary, sequence evolution, especially at the C/EBP and AP-1 motifs, directly affects the ability of LTR18A to act as regulatory elements, and most mutations have led to a decrease in regulatory potential.

### Evidence of selection for enhancer associated C/EBP and AP-1 motifs

Given that LTR18A has regulatory potential in certain cellular contexts like HepG2, we explored the possibility of host exaptation through the lens of selection. We first asked if LTR18A elements in chimpanzee, gorilla, gibbon, baboon, rhesus macaque, and marmoset have increased substitution rates compared to their human orthologs with respect to the distance between genomes. On average, LTR18A orthologs have slightly elevated substitution rates (12-32%) than the corresponding genomes (Supplemental Table 2). The increased substitution rate holds true even when only considering masked regions of the genome. Although it is possible that the genomic background rate includes regions under selection, the LTR18A substitution rates across primate species are overall inconsistent with purifying selection for the subfamily. Furthermore, both PhyloP and PhastCons scores at LTR18A elements provide no evidence of selection at the subfamily level across 30 mammals, including 27 primates^38, 39^ (Supplemental Figure 9).

While there is no evidence that LTR18A as a whole is under selection, it is possible that certain regions within LTR18A are. We aligned LTR18A elements in each of our seven primate species to the LTR18A consensus and tested sliding 10bp windows for increased conservation compared to the average window. Overall, 29% (707/2429) of all 10bp windows are significantly more conserved than the average window. The majority (84%) of conserved 10bp sliding windows are shared across all seven primates for a total of 24.5% (85/347) possible 10bp windows covering 58% of the LTR18A consensus (208/357bp) being classified as conserved. Shared, conserved regions defined by our sliding window analysis contain transcription factor motifs, including AP-1 and C/EBP (Figure 5A).

**Figure 5:**
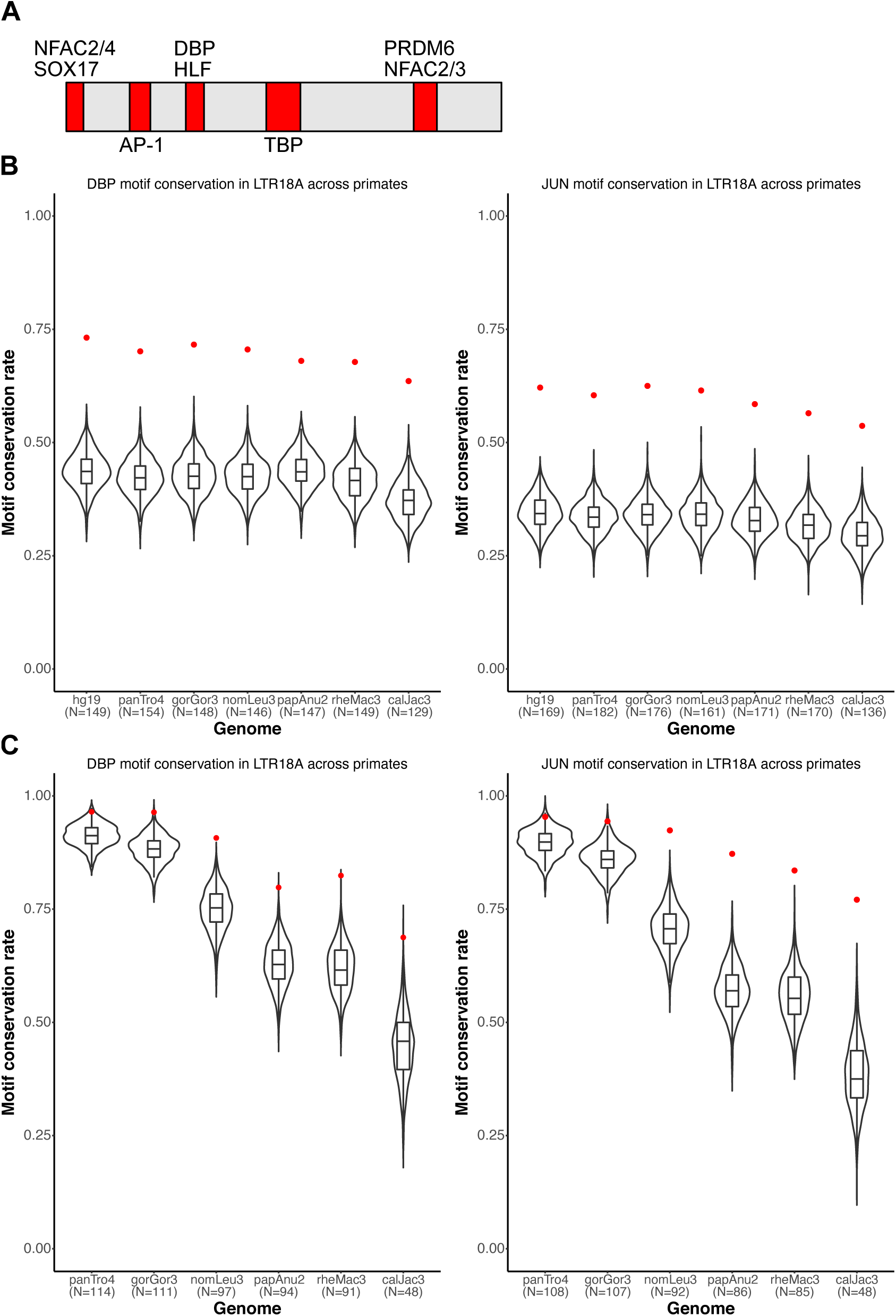
DBP and JUN motifs are more conserved than expected. A) Motifs that are fully encompassed within shared, conserved 10bp sliding windows across seven primate species. Motif locations in red are relative to the LTR18A RepBase consensus sequence. B) Distribution of expected neutral DBP and JUN motif conservation rates from the consensus motif. 1000 simulations are displayed for each species. The observed conservation rate is shown by the red point. C) Same as B, but for conservation rates from the hg19 ortholog as reference.

Since C/EBP and AP-1 motifs are critical for enhancer activity, we hypothesized that the motifs provided by LTR18A have been under selection and consequently exhibit higher conservation than expected under a neutral model of evolution. To obtain the background motif conservation rates, we adapted a method previously used in yeast^40^. Briefly, we take the sum of probabilities for all sequences that match a motif PWM, with each sequence probability calculated starting from the LTR18A consensus and the observed transition and transversion rate of the LTR18A subfamily. As in previous analyses, we chose DBP and JUN to represent C/EBP and AP-1. Expected conservation rates for DBP and JUN are consistent across species, ranging from 38.7% in marmoset to 44.8% in human for DBP and 34.1% in marmoset to 39.3% in human for JUN (Table 1). Meanwhile, observed DBP and JUN conservation rates are on average 69.3% and 59.3%, respectively, which is 26.4% and 21.6% higher than expected. This indicates that C/EBP and AP-1 motifs from the ancestral LTR18A sequence are being retained and may be under selection. Measuring conservation from the LTR18A consensus includes the transposition phase of TE evolution, which could select for C/EBP and AP-1 motifs due to enhancing transcription of the ERV. To address conservation specifically during primate evolution, we recalculated conservation rates by comparing human LTR18A elements to their primate orthologs. Generally, DBP and JUN motifs are significantly more conserved than expected (Table 2). The one exception is JUN for the human-chimpanzee comparison, which might be due to low human-chimpanzee divergence. We also confirmed higher motif conservation rates during transposition+speciation and speciation phases using simulations based on observed transition and transversion rates (Figure 5B, 5C). Together, our analysis suggests that C/EBP and AP-1 motifs contributed by LTR18A have been under selection in primates both before and after speciation.

**Table 1:** DBP and JUN motif conservation from RepBase consensus (ancestral), neutral evolution expectation vs. observed

| Motif: DBP_HUMAN.H11MO.0.B |  |  |  |  |  |  |
| --- | --- | --- | --- | --- | --- | --- |
| Species | Total possible elements | Expected conserved probability | Expected conserved number | Observed conserved number | Observed conserved proportion | p-value |
| hg19 | 149 | 44.77% | 66.71 | 109 | 73.15% | 1.61E-12 |
| panTro4 | 154 | 43.70% | 67.30 | 108 | 70.13% | 1.89E-11 |
| gorGor3 | 148 | 43.85% | 64.90 | 106 | 71.62% | 4.96E-12 |
| nomLeu3 | 146 | 44.10% | 64.39 | 103 | 70.55% | 6.12E-11 |
| papAnu2 | 147 | 42.94% | 63.12 | 100 | 68.03% | 3.97E-10 |
| rheMac3 | 149 | 42.17% | 62.84 | 101 | 67.79% | 1.22E-10 |
| calJac3 | 129 | 38.71% | 49.93 | 82 | 63.57% | 3.39E-09 |
| Motif: JUN_HUMAN.H11MO.0.A |  |  |  |  |  |  |
| Species | Total possible elements | Expected conserved probability | Expected conserved number | Observed conserved number | Observed conserved proportion | p-value |
| hg19 | 169 | 39.34% | 66.49 | 105 | 62.13% | 6.63E-10 |
| panTro4 | 182 | 38.54% | 70.14 | 110 | 60.44% | 6.33E-10 |
| gorGor3 | 176 | 38.65% | 68.02 | 110 | 62.50% | 4.05E-11 |
| nomLeu3 | 161 | 38.61% | 62.16 | 99 | 61.49% | 1.23E-09 |
| papAnu2 | 171 | 37.58% | 64.27 | 100 | 58.48% | 8.41E-09 |
| rheMac3 | 170 | 37.01% | 62.92 | 96 | 56.47% | 7.43E-08 |
| calJac3 | 136 | 34.07% | 46.33 | 73 | 53.68% | 7.01E-07 |

**Table 2:** DBP and JUN motif conservation from hg19 ortholog as reference, neutral evolution expectation vs. observed

| Motif: DBP_HUMAN.H11MO.0.B |  |  |  |  |  |  |
| --- | --- | --- | --- | --- | --- | --- |
| Species | Total possible elements | Expected conserved probability | Expected conserved number | Observed conserved number | Observed conserved proportion | p-value |
| panTro4 | 114 | 92.33% | 105.26 | 110 | 96.49% | 0.0476 |
| gorGor3 | 111 | 89.42% | 99.25 | 107 | 96.40% | 0.0084 |
| nomLeu3 | 97 | 76.83% | 74.53 | 88 | 90.72% | 0.0006 |
| papAnu2 | 94 | 65.84% | 61.89 | 75 | 79.79% | 0.0022 |
| rheMac3 | 91 | 64.71% | 58.89 | 75 | 82.42% | 0.0002 |
| calJac3 | 48 | 47.71% | 22.90 | 33 | 68.75% | 0.0018 |
| Motif: JUN_HUMAN.H11MO.0.A |  |  |  |  |  |  |
| Species | Total possible elements | Expected conserved probability | Expected conserved number | Observed conserved number | Observed conserved proportion | p-value |
| panTro4 | 108 | 91.08% | 98.37 | 103 | 95.37% | 0.0590 |
| gorGor3 | 107 | 87.70% | 93.84 | 101 | 94.39% | 0.0175 |
| nomLeu3 | 92 | 73.86% | 67.95 | 85 | 92.39% | 2.62E-05 |
| papAnu2 | 86 | 62.02% | 53.33 | 75 | 87.21% | 7.41E-07 |
| rheMac3 | 85 | 60.87% | 51.74 | 71 | 83.53% | 9.29E-06 |
| calJac3 | 48 | 44.93% | 21.57 | 37 | 77.08% | 3.77E-06 |

### Human LTR18A has epigenetic signatures of active regulatory elements

Our MPRA reveals that LTR18A elements have the sequence features to be activating regulatory elements depending on cellular context. To explore the relationship between regulatory potential from MPRA and enhancer function in the genome, we examined epigenetic marks in HepG2 and K562 using ENCODE data^41^. We first profiled LTR18A elements overlapping ATAC peaks for open chromatin, which is a common epigenetic feature for active regulatory elements. In HepG2, LTR18A is not enriched for ATAC peaks, with only 5 LTR18A elements overlapping with peaks. On the other hand, K562 has 11 overlapping LTR18A elements. This contrasts with the high MPRA activity in HepG2 relative to K562. Additionally, H3K27ac and H3K4me1, histone marks commonly associated with active enhancers, are also low across LTR18A in HepG2 and K562 (Supplemental Figure 10). We hypothesized that epigenetic repression of LTR18A may be the cause for the lack of active enhancer marks in HepG2. Consistent with this hypothesis, repressive histone mark H3K9me3 is enriched over LTR18A compared to the surrounding genomic region (Supplemental Figure 10). These results suggest that although LTR18A elements possess the sequence features necessary for enhancer activity, they can be epigenetically silenced.

While most of the LTR18A subfamily is unlikely to be active in HepG2 and K562, we sought to ascertain the contribution of LTR18A to the regulatory genome across human cell types and tissues. To get a global perspective, we overlapped LTR18A elements with candidate cis-regulatory elements (cCREs) as defined by ENCODE Registry V2 across 839 cell/tissue types^41^. Despite the limited number of cell/tissue types (25) that have full classification of cCREs, 69 of 198 (34.8%) LTR18A elements overlap with a cCRE, most of which (87%) have enhancer-like signatures (ELS) in at least one cell/tissue type. This represents 29.3% of all LTR18A bases which is about 3.1x enriched over the genomic background (p<3.5e-10, BEDTools fisher). Among fully classified cell/tissue types, keratinocytes have the highest number of LTR18A elements associated with ELS, followed by PC-3 and PC-9 cell lines (Figure 6A). LTR18A is not restricted to a single cell/tissue type, as some LTR18A elements are associated with cCREs in multiple cell/tissue types (Figure 6B). Across all 839 cell/tissue types, cell types with the most LTR18A overlapping cCREs largely consist of epithelial cells, such as MCF10A, mammary epithelial cells, esophagus epithelial cells, and foreskin keratinocytes (Figure 6C). To corroborate cCRE results which are based on DNase hypersensitivity, H3K27ac, H3K4me3, and CTCF ChIP-seq, LTR18A elements were intersected with ENCODE ATAC-seq peaks across 46 cell/tissue types. Similar to cCREs, LTR18A is especially enriched for ATAC peaks in epithelial cells/tissues foreskin keratinocytes and esophagus mucosa (11.4x and 16.1x enrichment over background respectively, BEDTools fisher). While certainly not comprehensive, the available epigenetic data supports an active enhancer-like state for LTR18A with the highest enrichment in epithelial cells.

**Figure 6:**
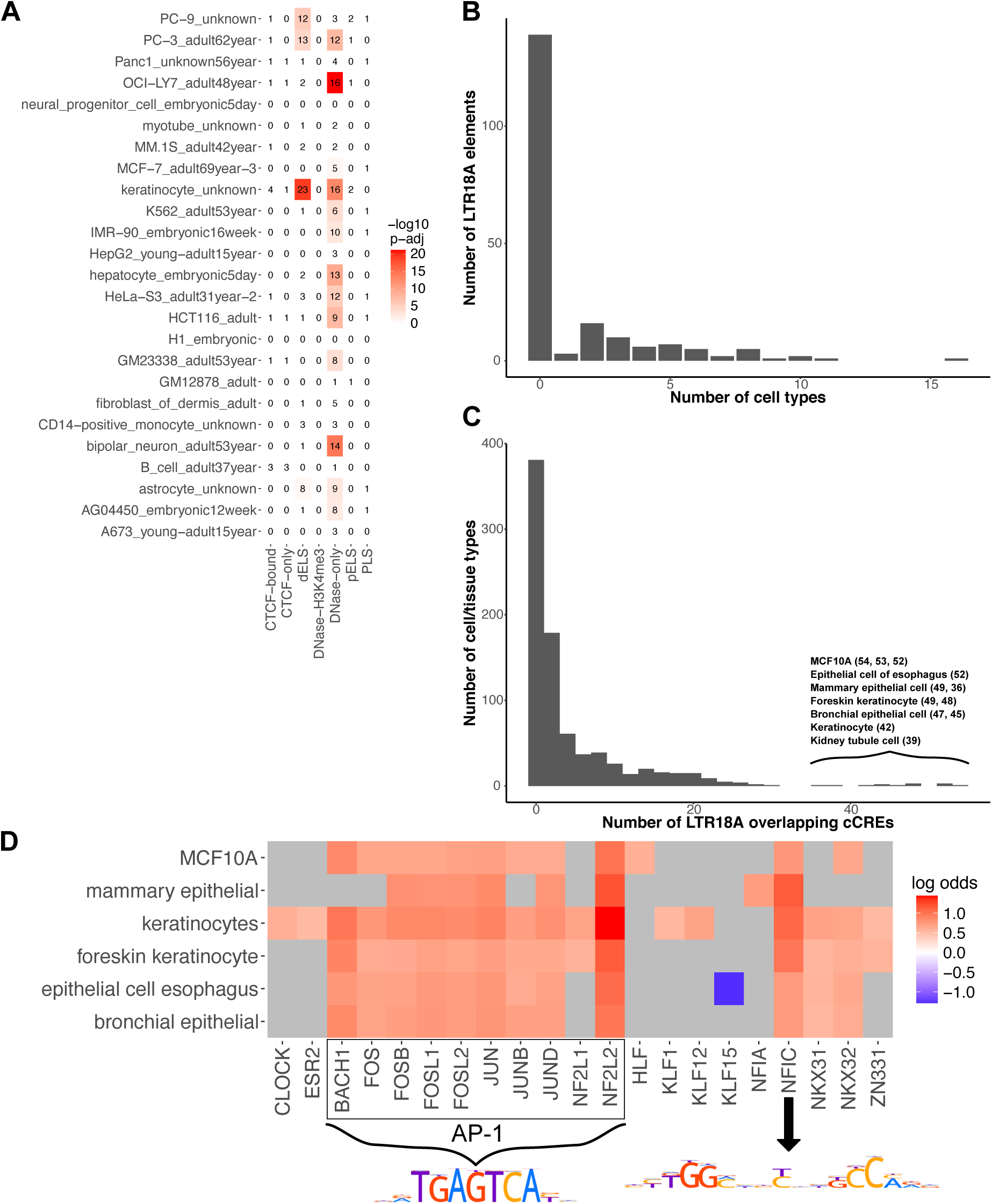
LTR18A elements are associated with enhancer epigenetic marks in human. A) Overlap of LTR18A with ENCODE cCREs across 25 full classification cell/tissue types (dELS, distal enhancer-like signature; pELS, proximal enhancer-like signature; PLS, promoter-like signature). The number of elements that overlap with cCREs are shown as well as their -log10 adjusted p-value by bedtools fisher. B) Distribution of LTR18A elements overlapping cCREs across multiple full classification cell/tissue types. C) Distribution of cell/tissue types overlapping LTR18A elements. The top cell/tissue types are displayed with the number of LTR18A elements that overlap with a cCRE. D) Motifs associated with the cCRE-overlapping LTR18A elements from the top cell/tissue types in C. Grey indicates non-significance at adjusted p-value threshold of 0.05. PWMs for JUN (AP-1 related factors) and NFIC are shown.

As LTR18A enhancer potential is influenced by sequence variation especially at transcription factor binding sites, we sought to understand whether transcription factor motifs are associated with active epigenetic states. Without considering cell/tissue type, we found no transcription factor motif to be significantly associated with LTR18A overlapping cCREs relative to other LTR18A. Due to the cell type specific nature of most enhancers, we identified motifs enriched in cCRE associated LTR18A in the top cell/tissue types (Figure 6D). Many of the most common motifs are of AP-1 transcription factors. Another common motif is NFIC, which is consistent with an activating role previously described in cancer and could serve a similar role in activating LTR18A elements^42^. Of note, the C/EBP-related factor HLF is enriched only in the MCF10A cell line. Using ATAC data, we confirmed AP-1 and NFIC motifs as enriched in LTR18A elements associated with active epigenetic states in foreskin keratinocytes and esophagus mucosa. Altogether, these results suggest that LTR18A elements become epigenetically activated in epithelial cells primarily through AP-1 transcription factors and NFIC.

## Discussion

Since Britten and Davidson first hypothesized how repetitive elements could influence the development of gene regulatory networks, a growing number of studies have shown the contribution of TEs as regulatory modules^43^. Using LTR18A as a representative subfamily, we performed the first systematic functional testing of regulatory potential for a TE subfamily using MPRA. By taking advantage of the natural sequence variation across elements, we identify AP-1 and C/EBP-related motifs as important drivers of LTR18A regulatory activity. This regulatory activity is highly dependent on cell context, with LTR18A displaying much higher activity in HepG2 than in K562. However, the sequence potential for regulatory activity does not necessarily reflect activity in the genome, as shown by LTR18A elements rarely associating with active epigenetic marks in HepG2. Due to general repression of TEs, we believe that similarly silenced TEs with the potential for enhancer activity may be common. These inactive TEs may be latent under epigenetic control, but there remains the possibility that a changing epigenome such as during tumorigenesis can reactivate them^44^.

Another unique aspect of this study is leveraging the phylogenetic relationship between LTR18A elements within human and across primate species to investigate the origin and evolution of regulatory activity in the subfamily. Previous research has implicated two evolutionary paths through which TE sequence can contribute to the spread of regulatory modules. The first case is when the ancestral TE originally possesses the driving regulatory features, such as the p53 binding site in LTR10 and MER61 or the STAT1 binding site in MER41B^20, 26^. A second possibility exists where the ancestral TE gains the regulatory module in one lineage through mutation before amplification, such as the 10bp deletion in ISX relative to ISY in *D. miranda* that recruits the MSL-complex^45^. In the LTR18A family, we observe both scenarios. Both C/EBP and AP-1 motifs are found in the LTR18A consensus and our reconstructed subfamily ancestor, and many elements retain the motifs to the present day. Divergence from the ancestor over time, especially at the two motifs, is correlated with a decrease in regulatory activity. In addition to the two consensus motifs, a second C/EBP motif is gained through an insertion at an early evolutionary timepoint. This second C/EBP motif further increases the regulatory potential of LTR18A. Ultimately, however, few present-day elements have maintained the second motif. This could be explained by negative selection or a deletion bias from the sequence similarity of the insertion with the upstream sequence. It also plausible that our evolutionary reconstruction makes an incorrect assumption about the timing of the second C/EBP motif, and each one occurred independently rather than through a common ancestor. If this scenario is true, recurrent insertions in TEs may be more common than previously thought.

An intriguing possibility is the relationship between TE regulatory potential and genomic expansion. In our reconstructed LTR18A phylogenetic tree, we observe loss of enhancer activity in the leftmost lineage going as far back as its lineage ancestor. This low enhancer activity lineage corresponds to the earliest diverging branch in the human LTR18A subfamily phylogenetic tree and composes only ∼1/6 of all elements. On the other hand, the major lineage of LTR18A has enhancer activity throughout transposition. The stark contrast between the two lineages in enhancer activity and abundance leads us to speculate that the regulatory potential of LTR18A was directly related to its ability to expand in the genome. This is perhaps unsurprising, as transcription is typically the first step of transposition and provides the substrate for integration of retrotransposons. However, one important consequence is that transcription factor binding sites that contribute to TE regulatory potential could be enriched within a subfamily due to biased lineage amplification. This appears to have been the case for the recently reclassified LTR7 subfamilies, each of which possess a unique set of transcription factor motifs and underwent a wave of genomic expansion to fill different early embryonic niches^46^. It will be important for future studies to distinguish between selection and passive enrichment of transcription factor binding sites through lineage amplification.

To compare ancestral and present day LTR18A elements, we tested all elements within the same cell line. This assumes that HepG2 and K562 cells provide the same *trans* environment as the equivalent primate and ancestral cell types. Previous studies suggest that transcription factor binding and subsequent activation of transcription are deeply conserved from humans to flies^47, 48^. Klein et al. make a similar assumption in their study of liver enhancer evolution in primates and find the same general trend that present-day elements have lost enhancer activity relative to the ancestral state^18^.

Most TEs are thought be under neutral evolution and do not significantly impact phenotype. We find that LTR18A elements as a whole have higher mutation rates than genomic average and do not exhibit signs of selection based on phyloP and phastCons scores. Despite the lack of evidence for selection at the element level, AP-1 and C/EBP binding motifs found within LTR18A are more conserved than expected under the neutral model of evolution. This suggests that selection does not need to apply to entire TEs and instead acts on functional units found within each element. Indeed, we find that at least a third of LTR18A elements have enhancer associated epigenetic marks, and in some cell/tissue types, the active elements are enriched for the conserved AP-1 motif. Although the C/EBP motif is not significantly enriched with active elements outside of MCF10A, we suspect that the motif is important in other cell/tissue types that have yet to be profiled.

## Methods

### LTR18A manual curation for ancestral reconstruction

We downloaded RepeatMasker 4.0.5 (Repeat Library 20140131) annotations for human (hg19), chimpanzee (panTro4), gorilla (gorGor3), gibbon (nomLeu3), rhesus macaque (rheMac3), and marmoset (calJac3) genomes^49^. For baboon (papAnu2) which is not available on www.repeatmasker.org, we ran RepeatMasker 4.1.0 using the RepBase RepeatMasker library 20170127. Since LTR18A consensus sequences are 98% similar between the two repeat libraries, we believe that most if not all LTR18A elements will be identified in papAnu2 in the same way as the other primate genomes. For the closest two subfamilies, LTR18B and LTR18C consensus sequences are ∼75% and 67% similar to the LTR18A consensus respectively.

For manual curation, we examined the alignment of each annotated LTR18A element and removed the element if it satisfied any of our filtering criteria (Supplemental Table 1). First, we exclude LTR18A elements that have significant alignments to LTR18B or LTR18C. RepeatMasker outputs alignment scores for each repetitive element, some of which have multiple significant alignment scores for different subfamily consensus sequences. RepeatMasker then chooses the subfamily with the highest alignment score to annotate elements with the same ID. A consequence of this method is that fragmented elements can be annotated for the same subfamily even when the highest scoring alignment differs for each fragment. Since LTR18B and LTR18C consensus sequences are ∼75% and 67% similar to LTR18A respectively, some LTR18A elements have significant alignments to LTR18B and/or LTR18C. Thus, we discard these elements with multiple possible alignments to avoid ambiguity from subfamily assignment. Second, we use paired LTR information to remove LTR18A elements that have discordant annotations. Due to the mechanism of ERV retrotransposition, we expect non-solo LTR18A elements to exist as same orientation pairs that are separated by the ERV internal region. Using this logic, we reasoned that paired LTRs that are assigned to different subfamilies have uncertain annotation.

To find LTR18A ortholog sets for ancestral reconstruction, we searched for LTR18A element pairs that fulfilled several requirements. First, the hg19 LTR18A elements must have orthologs in chimpanzee and gorilla. Second, elements must have orthologs in at least two of the other primate species: gibbon, baboon, rhesus macaque, and marmoset. Third, hg19 LTR18A elements must be >250bp (>70% of consensus) in length. Finally, both elements of a pair need to pass all requirements to be selected for ancestral reconstruction. Orthologs were defined using the chain files from UCSC to find LTR18A elements within the same syntenic blocks^31^.

Ancestral reconstruction of both ortholog ancestors and subfamily ancestors used MAFFT and PRANK followed by manual curation^32, 33^. To generate ortholog ancestors, we aligned ortholog sets (e.g. human, chimpanzee, gorilla, gibbon, baboon orthologs) using MAFFT multiple sequence alignment. We used the alignments to produce ancestral and intermediate sequences as well as the phylogenetic tree using PRANK. The PRANK phylogenetic trees typically reflected the expected evolutionary relationship between the seven primate species. Next, we manually adjusted ortholog ancestors to remove unlikely insertions. We focused on insertions rather than deletions due to the possibility of insertions propagating up the tree. We determined insertion sites by examining the multiple sequence alignment of ortholog ancestors and finding gaps in the alignment created by insertions in only a few ortholog ancestors. Generally, we used parsimony when deciding to keep or remove an insertion. For example, if the insertion is present in only one primate lineage, then it is less likely for the insertion to have existed in the ortholog ancestor. Our reasoning is that an insertion in the ortholog ancestor and subsequent deletion in the other lineages requires at least two mutation events, whereas a single insertion in one primate lineage requires only one mutation event. After manual curation of ortholog ancestors, we used MAFFT and PRANK to reconstruct the phylogenetic tree and sequences of LTR18A subfamily ancestral sequences. We again applied parsimony to manually adjust the LTR18A subfamily ancestor.

### LTR18A MPRA library construction

The MPRA library was designed to consist of a motif-focused half and a tiling half. To design the motif-focused half of our MPRA library, we took advantage of the relatedness of TEs within the same subfamily. Similar to RepeatMasker, we can align all LTR18A elements to a reference sequence. Instead of using the subfamily consensus sequence, we used our reconstructed ancestral node 43 to perform pairwise global alignments to all present-day and reconstructed elements. Then, we took the sequence of each element aligned to the first 160bp of ancestral node 43. We filtered out elements that have fewer than 70bp due to deletions and elements that have more than 160bp due to insertions. We also removed elements that contain a restriction site that we used for cloning. In total, 1255/1387 RepeatMasker annotated LTR18A elements across seven primate genomes and all 280 reconstructed elements were included. For the tiling half of the library, we selected all pre-speciation ancestral reconstruction elements, ortholog ancestors and their present-day hg19 elements, and the LTR18A consensus. We then synthesized 160bp tiles at 10bp intervals spanning each selected element. In addition to motif-focused and tiled sequences, we selected 456 elements for reverse complements, 37 pairs of elements for JUN mutagenesis, and 46 pairs of elements for DBP mutagenesis. Elements for mutagenesis were chosen based on the closest primate ortholog with/without the motif. JUN motifs were mutated to TCACCAATGGT and DBP motifs were mutated to TCCCACAGCAT. Non-motif containing elements were mutated to GCTGAGTCATG for JUN and ATTATGTAACC for DBP. For positive and negative controls, we selected 223 regions from a previous study by Ernst et al.^12^. 30 dinucleotide shuffled LTR18A RepBase consensus sequences were included as a second set of negative controls^50^. Each synthesized sequence was tagged with 10 unique barcodes. To control for differences in overall library activity between cell lines, we included a set of sequences that would leave only the basal hsp68 promoter tagged with 300 barcodes. Oligos were ordered from Agilent and structured as follows: 5’ priming sequence containing NheI site (CGGTATCTAAGAgctagcGT)/CRE/EcoRI site/Filler (if necessary)/BglII site/BamHI site/constant ‘G’/barcode/constant ‘A’/AgeI site/3’ priming site (ATTAGCATGTCGTG)^11^. Total length of oligos was 230bp. In total, 5918 elements were synthesized using 59470 unique barcodes.

The MPRA library was constructed as previously described with some adjustments. An AgeI site was introduced upstream of the SV40 polyA signal and the BamHI site downstream of the SV40 polyA signal was deleted using the QuikChange Lightning site-directed mutagenesis kit (Agilent). Synthesized oligos were amplified with 0.05pmol of template per 50μL PCR reaction for seven cycles using MPRA library amplification primers. A total of 32 reactions were performed. Following amplification and gel purification, oligos were cloned into a pGL backbone with the AgeI insert using NheI and AgeI sites. Multiple ligations were pooled, purified by PCR cleanup (Nucleospin), and transformed into 5-alpha electrocompetent *E. coli* (NEB). The hsp68 promoter driving dsRed reporter was cloned using EcoRI and BamHI sites. The MPRA library with the hsp68 promoter and dsRed reporter was purified and transformed into *E. coli* before plasmid extraction. The final library was concentrated by ethanol precipitation.

### Cell culture and library transfection

Cell culture and library transfections were performed as previously described^11^. K562 cells were grown in RPMI 1640 with L-glutamine (Gibco) + 10% Fetal Bovine Serum (FBS) + 1% penicillin/streptomycin. HepG2 cells were grown in Dulbecco’s Modified Eagle Medium with high glucose, L-glutamine, and without sodium pyruvate + 10% FBS + 1% penicillin/streptomycin. For each of three replicates, 5 μg of library was transfected into 1.2 million cells using Neon electroporation (Life Technologies). For K562, electroporation parameters were three 10 millisecond pulses at 1450V. For HepG2, electroporation parameters were three 20 millisecond pulses at 1230V. As a transfection control, 0.5 μg of pmaxGFP (Lonza) was used.

### Measurement of library expression

RNA extraction was performed 24 hours after transfection using PureLink RNA Mini Kit with on-column DNase treatment (Life Technologies) followed by DNase I treatment using TURBO DNA-free kit (Invitrogen). Samples were prepared for RNA-seq as previously described^11^. First strand cDNA synthesis was performed using Superscript III Reverse Transcriptase (Life Technologies). Barcodes were amplified from cDNA from three transfections and three technical replicates of DNA from the plasmid library. Amplified barcodes were digested with KpnI and EcoRI and ligated to Illumina adapters. Ligation products were further amplified, after which replicates and plasmid library DNA input were pooled for sequencing. We obtained over 1000x average coverage for each transfection replicate and the DNA input. For each tested element, we added up read counts for all of its barcodes and filtered out those with fewer than 5 total counts in any transfection replicate or DNA input. Reads were then normalized to counts per million (CPM). Expression of an element was calculated as RNA CPM/DNA CPM. Expression was normalized to the average of Basal construct transfection replicates. Finally, enrichment score was calculated as the log2 of normalized expression. Enrichment scores of elements were highly reproducible across transfection replicates in HepG2 (average R^2^=0.904) while moderately reproducible in K562 (average R^2^=0.666) (Supplemental Figure 3). We confirmed that orientation does not have large effects on enrichment score in both HepG2 and K562 (Supplemental Figure 4). We also found that selected control sequences from Ernst et al. follow expected trends for both their original annotations as well as redefined annotations based on expression values from Ernst et al. MPRA results (Supplemental Figure 5). Enrichment scores of elements are provided in Supplemental Data.

### TE-WAS analysis of nucleotides and motifs

LTR18A sequences were first globally aligned pairwise to the ancestral node 43 sequence as reference^51^. Individual pairwise alignments were then combined based on the common reference. Positions that had bases (not gaps) in less than 20% of all LTR18A sequences were removed. This filter retained all consensus base positions.

GWAS analysis tool PLINK was used to identify nucleotides significantly associated with the phenotype, such as MPRA activity/inactivity or ATAC peak^36^. We limited tested nucleotides at each position to the most common nucleotide at the position across LTR18A sequences to give us greater confidence based on sample size. We ran PLINK association analysis using the above-described alignment and MPRA active/inactive annotations for each element based on enrichment score. Nucleotides were deemed significant if p-value < 5e-5.

From the list of significant nucleotides in TE-WAS, we identified transcription factor motifs that are overrepresented based on information content. Information content at each significant nucleotide was calculated from each motif’s position frequency matrix with the background nucleotide frequencies of 0.25. The information content of significant nucleotides within each motif was then compared to a background expectation derived from 1000 random shuffles of significant nucleotides for the phenotype. Motifs were identified if they had higher information content from significant nucleotides than background using t-test and more than significant nucleotide within the motif.

### Evolutionary analysis using SHARPR

From tiled MPRA, we calculated regulatory activity for full length elements using SHARPR with a few adjustments^12^. For each tile of an element, the previously calculated enrichment score was used as input for SHARPR infer with the default varpriors of 1 and 50. Each inferred 10bp step was then normalized to the mean inferred value for randomly shuffled Basal elements as background. SHARPR combine and interpolate commands were used to generate the SHARPR nucleotide activity scores. Finally, full length element activities were calculated as the sum of nucleotide scores across each element.

To validate the SHARPR approach, we identified motifs that were enriched in peaks, or regions of high nucleotide activity. Peaks were defined as regions with nucleotide activity scores greater than three standard deviations above the Basal mean. Enriched motifs were then identified in peak regions using AME using shuffled sequence as background^34^.

### Transcription factor motif conservation

For sliding window conservation analysis, we aligned all present-day genomic LTR18A elements to the RepBase consensus sequence using the previously defined method. Conservation, defined as percent match to the consensus, was calculated for each 10bp window for each element in each species. Windows with gaps or degenerate bases in at least half of the total window length (>=5) were excluded. The mean conservation was then calculated for each 10bp window separately for each species. Windows were determined to be significantly conserved using t-test comparing conservation across elements in the window against conservation across all windows, with a p-value threshold of 0.05 after Bonferroni correction. Only windows that were conserved in all seven primate species were kept for further analysis. Motif scanning by FIMO was performed to find transcription factor motifs fully within conserved windows^37^.

For JUN and DBP transcription factor motif conservation analysis, transition and transversion rates in the LTR18A subfamily were calculated for each species. The neutral expectation for motif conservation was calculated as previously described^40^. We identified all kmers of the motif length which are found by FIMO^37^. The total motif conservation probability was calculated as the sum of the probabilities for each motif kmer. We used the RepBase consensus sequence as the ancestral LTR18A state. To represent post-speciation conservation, we used hg19 orthologs as the reference to compare to other primate LTR18A elements. The observed motif conservation rate was calculated for each species based on the percentage of elements that retain the motif. Elements with gaps in the alignment to its reference were excluded. Statistical significance was determined by one sample test of proportions and a p-value threshold of 0.05. We also simulated transcription factor motif conservation rates for each primate species. Each simulation consisted of randomly mutating nucleotides in the motif region of each LTR18A element based on the observed transition and transversion rates. 1000 simulations were performed for each motif.

### Overlap of LTR18A with genomic annotations

The cCRE genome annotations and various epigenetic datasets such as ATAC-seq, histone ChIP-seq, and WGBS were downloaded from ENCODE^41^. The phyloP and phastCons scores were downloaded from ENCODE and converted to bedGraph^31^. Overlaps with LTR18A elements were obtained by BEDTools intersect with the criteria of at least 50% LTR18A length overlapping with a cCRE or epigenetic mark peak^52^. Enrichment of LTR18A in cCREs and ATAC peaks was obtained by BEDTools fisher using the same criteria. Heatmaps at and around LTR18A were generated using deepTools^53^.

### Identification of motifs associated with cCRE overlapping LTR18A

Fisher’s exact test was used to determine if transcription factor binding motifs in LTR18A elements are associated with cCRE overlap. Motifs that had adjusted p-values below 0.05 were considered significant. The top six cell/tissue types were selected for analysis as they provided the greatest number of LTR18A elements overlapping cCREs.

### Subfamily age estimate

The average divergence, weighted by copy length, was calculated for the LTR18A subfamily using the RepeatMasker output for hg19. The age was obtained by using the average divergence and the average mammalian genome mutation rate of 2.2 x 10^-9^ per base per year^54^.

## Acknowledgements

We would like to thank J. Hoisington-López and M.L. Jaeger from The Edison Family Center for Genome Sciences & Systems Biology (CGSSB) for assistance with sequencing. This work was funded by NIH grant numbers R01HG007175, U01CA200060, U01HG009391, U41HG010972, and U24HG012070. A.Y.D. was supported by NHGRI training grant T32 HG000045.

## Contributions

A.Y.D., V.S., and T.W. designed the study. X.Z. contributed to evolutionary analysis. N.O.J. and N.L.S. contributed to TE-WAS analysis. A.Y.D. performed the MPRA with contributions by H.G.C. and B.A.C. in design and analysis. The manuscript was prepared by A.Y.D. and T.W. with input from authors.

