## Supplemental_material for "Evolution of transposable element-derived enhancer activity"

| Coordinate (hg19) | Reason for removal |
|---|---|
| chr2:53717712-53717918 | Alignment is to LTR18B |
| chr2:53717983-53718184 | Same element ID as chr2:53717712-53717918 (above) and LTR18A alignment truncated (1-210) |
| chr4:28012466-28012685 | Alignment is to LTR18B |
| chr4:28012746-28012949 | Same element ID as chr4:28012466-28012685 (above) and LTR18A alignment truncated (1-211), also paired LTR with LTR18C (chr4:28016239-28016607) |
| chr4:90025654-90025852 | Same element ID as chr4:90025918-90026134 (below) and LTR18A alignment truncated (1-211) |
| chr4:90025918-90026134 | Alignment is to LTR18B and LTR18A, alignment to LTR18B is used in rmsk out because it is longer |
| chr4:90029346-90029548 | Same element ID as chr4:90029626-90029704 and chr4:90029757-90029861 (both below), LTR18A alignment truncated (1-211) |
| chr4:90029626-90029704 | Alignment is to LTR18B |
| chr4:90029757-90029861 | Alignment is to LTR18B |
| chr10:59575924-59576130 | Alignment is to LTR18B and LTR18A, LTR18A alignment is truncated (285-357) but alignment to LTR18B is used in rmsk out because it is longer |
| chr11:21832342-21832596 | Alignment is to LTR18B (2 pieces) and LTR18A, LTR18A alignment is truncated (1-181) but alignment to LTR18B (combined 2 pieces) is used in rmsk out because it is longer |
| chr11:21832616-21832829 | Alignment is to LTR18B, also paired LTR with LTR18C (chr11:21835813-21836035) |
| chr1:191578310-191578437 | Paired LTR with LTR18B (chr1:191577527-191578089) |
| chr11:101626917-101627207 | Alignment is to LTR18B, LTR18C, and LTR18A (which is highest scoring), LTR18A alignment is truncated (1-210) but alignment to LTR18B is used in rmsk out because it is longer |
| chr11:101627272-101627424 | Alignment is to LTR18B and LTR18A, LTR18A alignment is truncated (245-355) but alignment to LTR18B is used in rmsk out because it is longer |
| chr18:22188578-22188720 | Alignment is to LTR18C and LTR18A, LTR18A alignment is truncated (225-357) but start is from LTR18C alignment while stop is from LTR18A alignment |
| chr18:51468362-51468410 | Alignment is to LTR18C |

Supplemental Table 1: Manual curation of hg19 LTR18A elements. Elements that were removed from ancestral reconstruction consideration are listed.

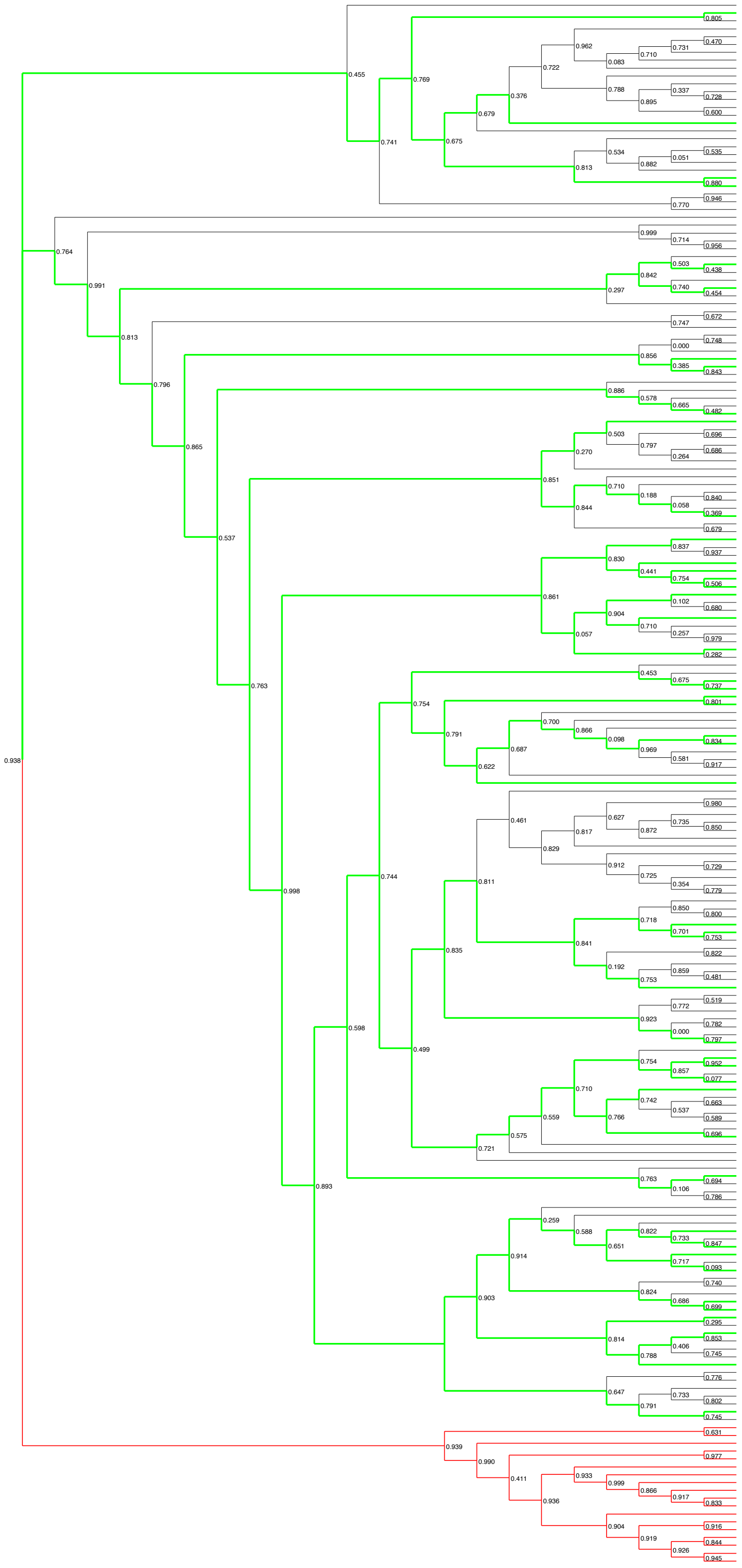

Supplemental Figure 1: Phylogenetic tree of manual curated hg19 LTR18A. Green lines indicate paths to hg19 LTR18A used in ancestral reconstruction. Red lines indicate paths to LTR18B elements used as an outgroup. Node labels are bootstrap values from FastTree.

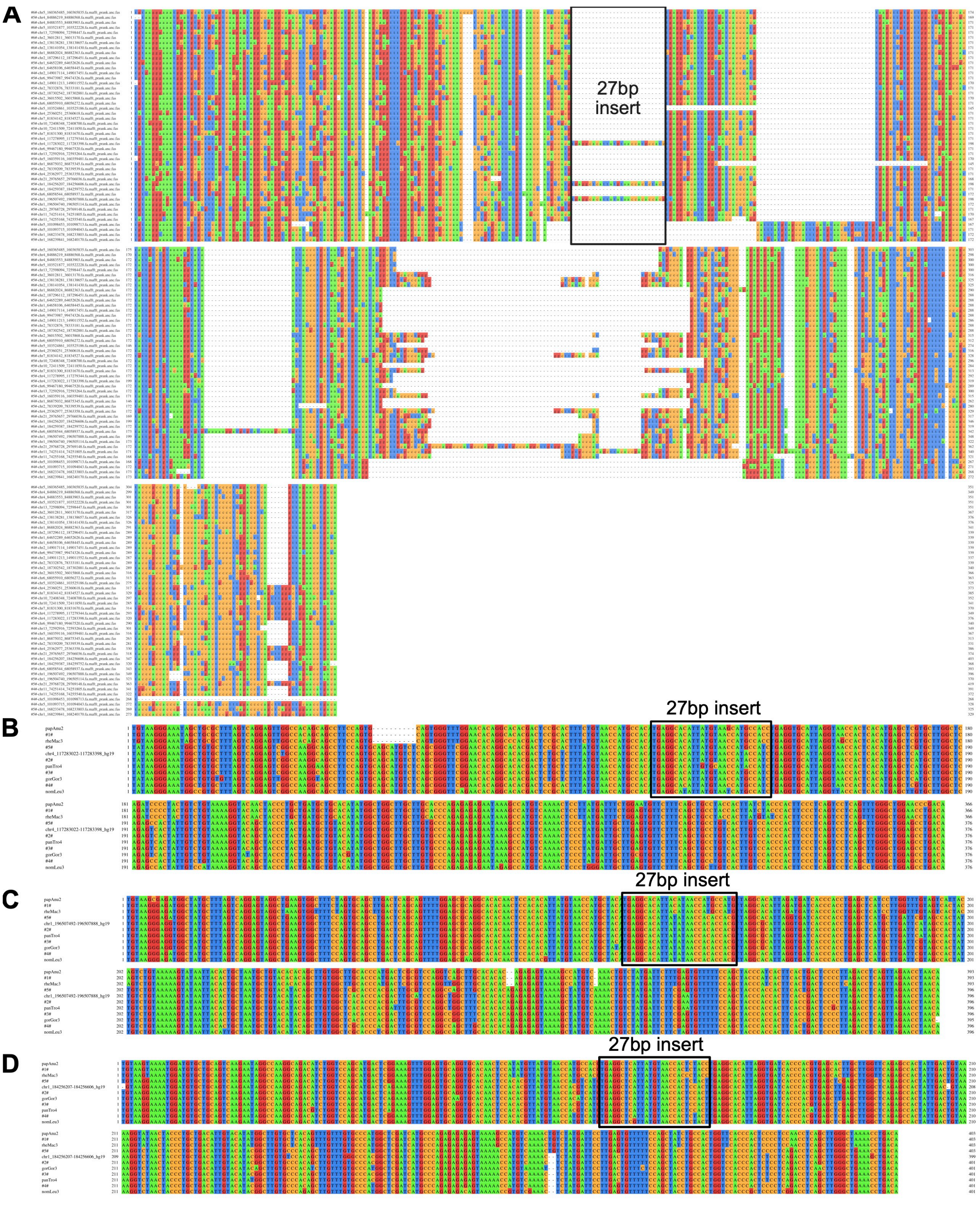


Supplemental Figure 2: MAFFT sequence alignment of LTR18A elements containing the 27bp insert. A) Alignment of all ortholog ancestors from the first round of ancestral reconstruction. The region of the 27bp insertion is boxed. B, C, and D) Alignments of the three sets of primate orthologs and their reconstruction intermediates that contain the 27bp insertion (boxed for each alignment). Coordinates for the hg19 reference element are displayed.

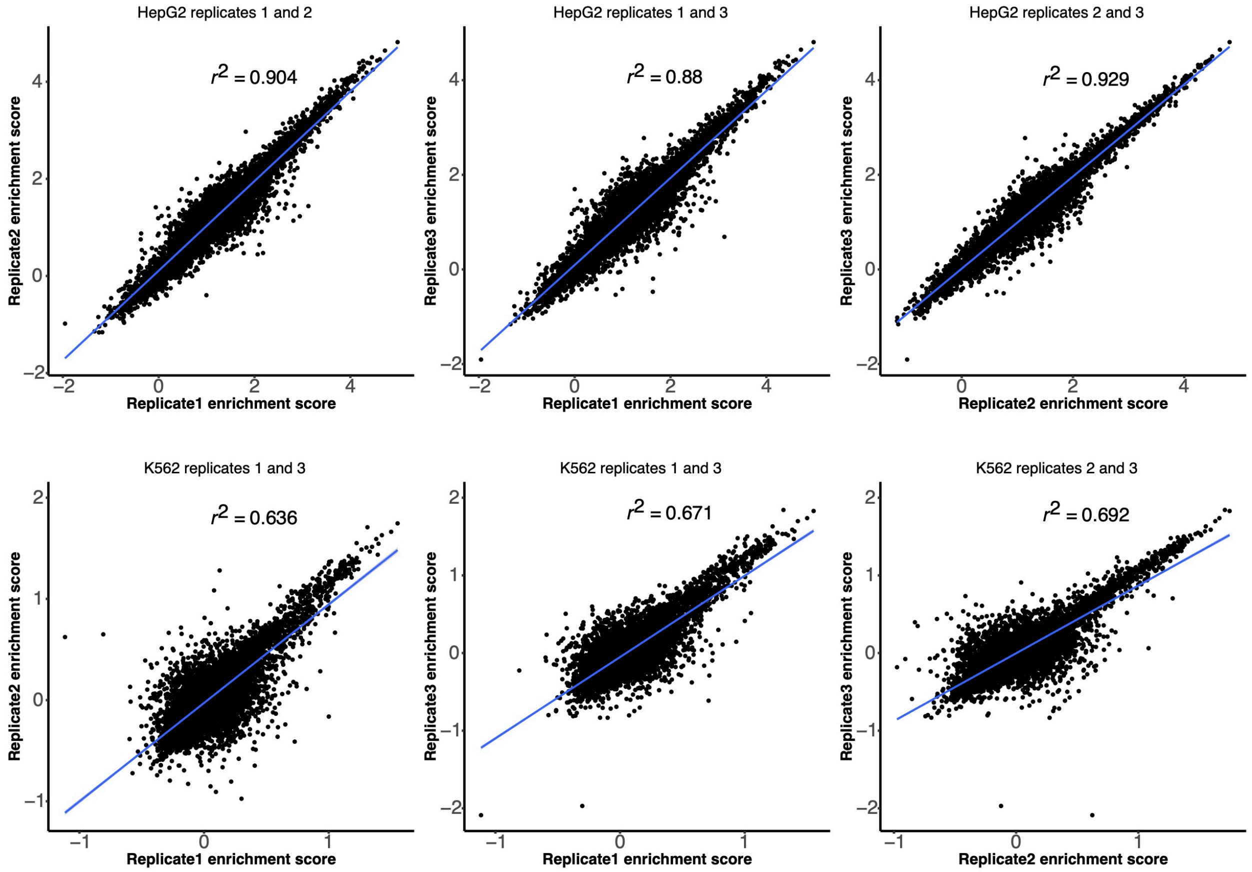


Supplemental Figure 3: Biological replicate enrichment score correlations for HepG2 and K562.

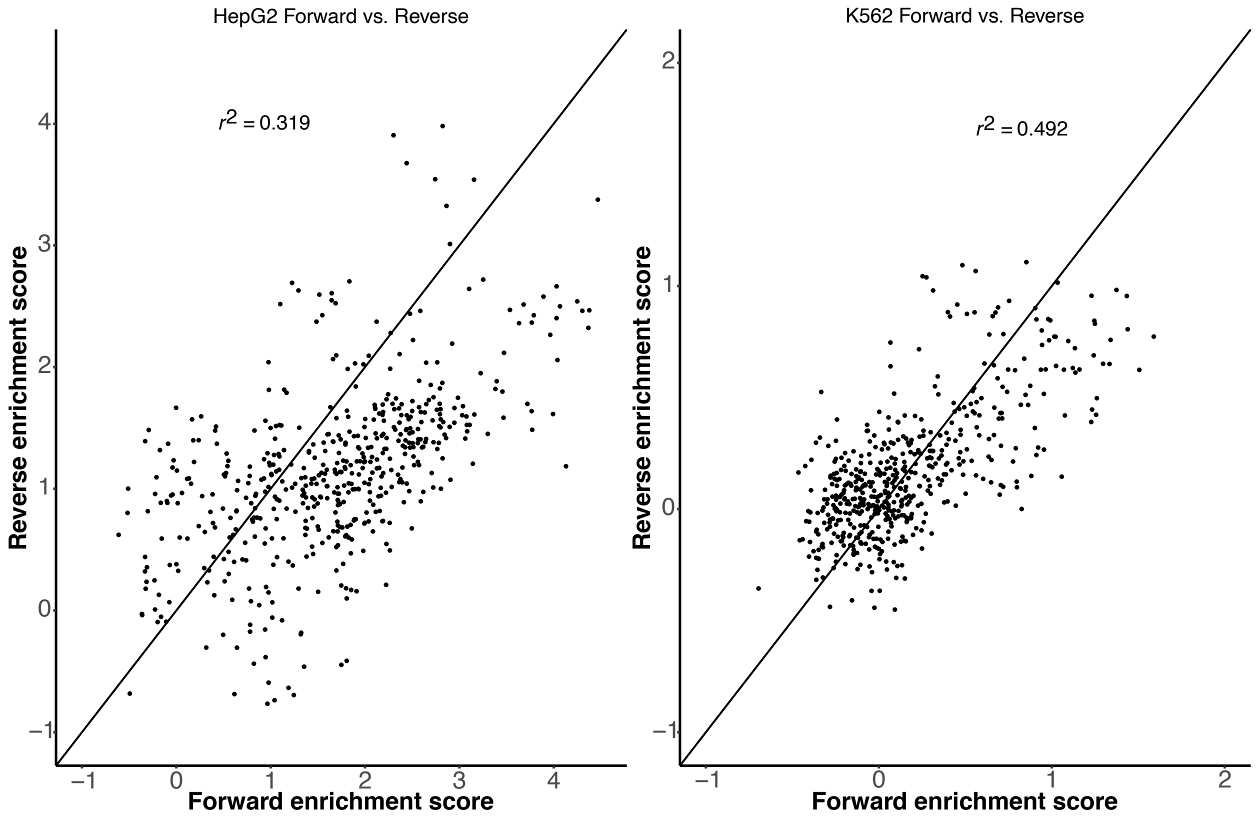


Supplemental Figure 4: Correlation of LTR18A fragments tested in forward and reverse orientations in HepG2 and K562.

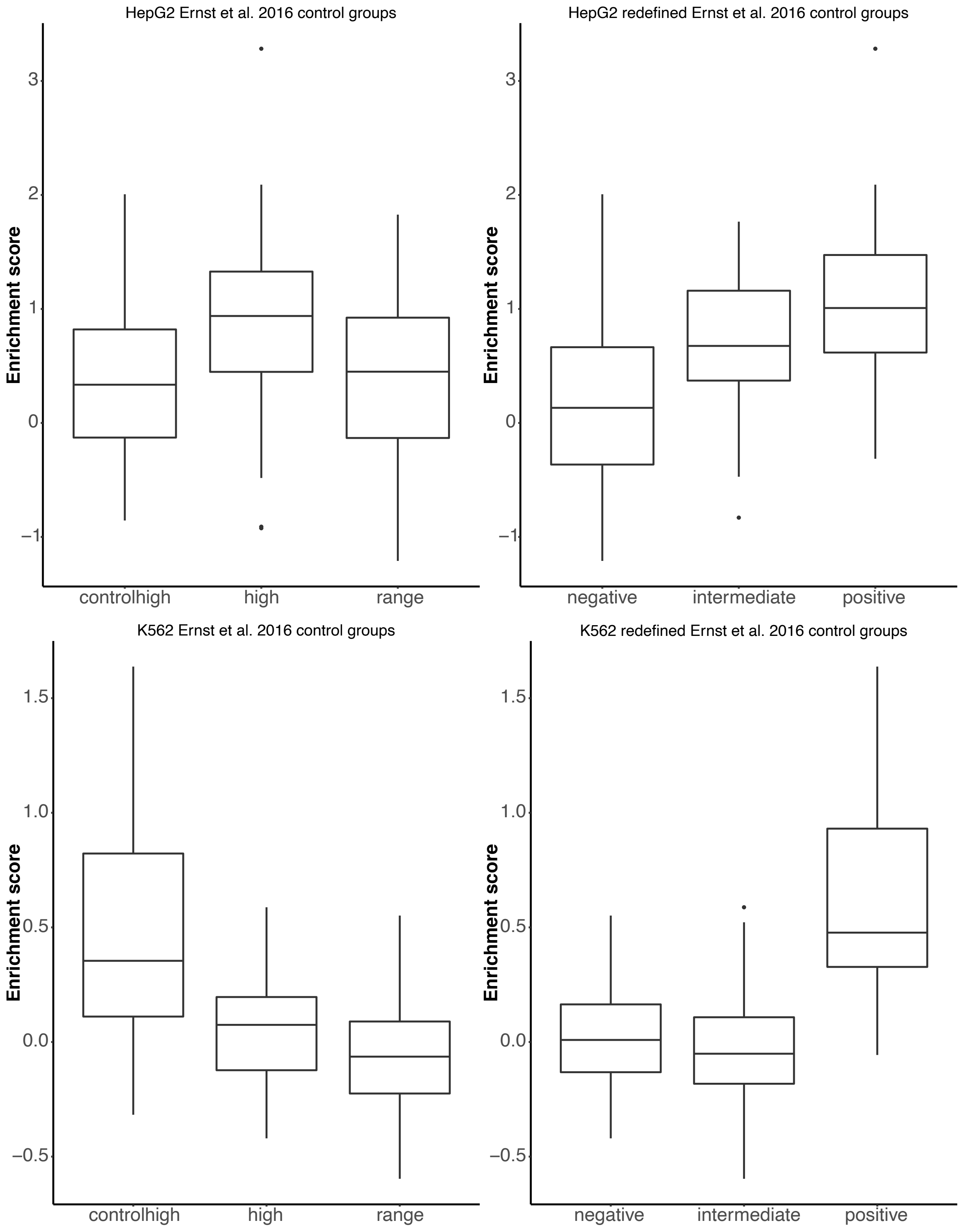


Supplemental Figure 5: Enrichment scores of controls from Ernst et al. 2016 in HepG2 and K562. Previously defined categories of sequences are as follows: "controlhigh" denotes candidate enhancer regions with strong H3K27ac dip scores in K562 but low activity states in

HepG2, “high” denotes candidate enhancer states in HepG2 with the strongest H3K27ac dip scores, “range” denotes regions with a range of dip scores. These same sequences were redefined to negative, intermediate, and positive groups based on their expression level in the Ernst et al. Pilot MPRA.

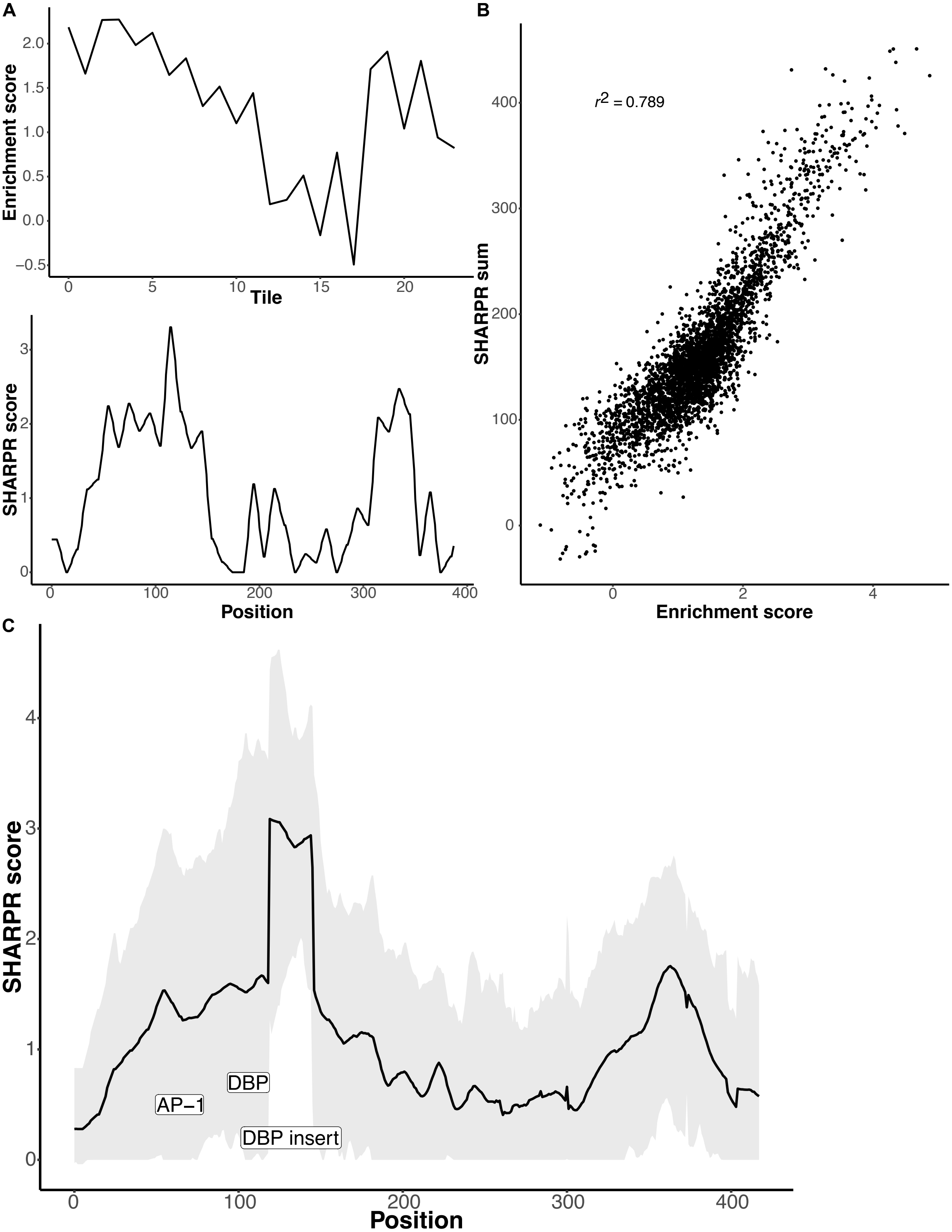

Supplemental Figure 6: SHARPR describes regulatory activity from tiling data. A) Transformation of tile enrichment scores (upper panel) to SHARPR nucleotide activity scores (lower panel) for the LTR18A subfamily reconstructed ancestor. B) Correlation of SHARPR score sums with enrichment scores for every tile. C) SHARPR scores across tiled LTR18A elements. Elements were aligned to obtain universal positions. The mean SHARPR score for each position is plotted in the solid black line, with 90% of all scores falling within the grey shaded region. AP-1 and DBP motif positions are labeled, with the DBP motif in the 27bp insert differentiated from the consensus DBP motif.

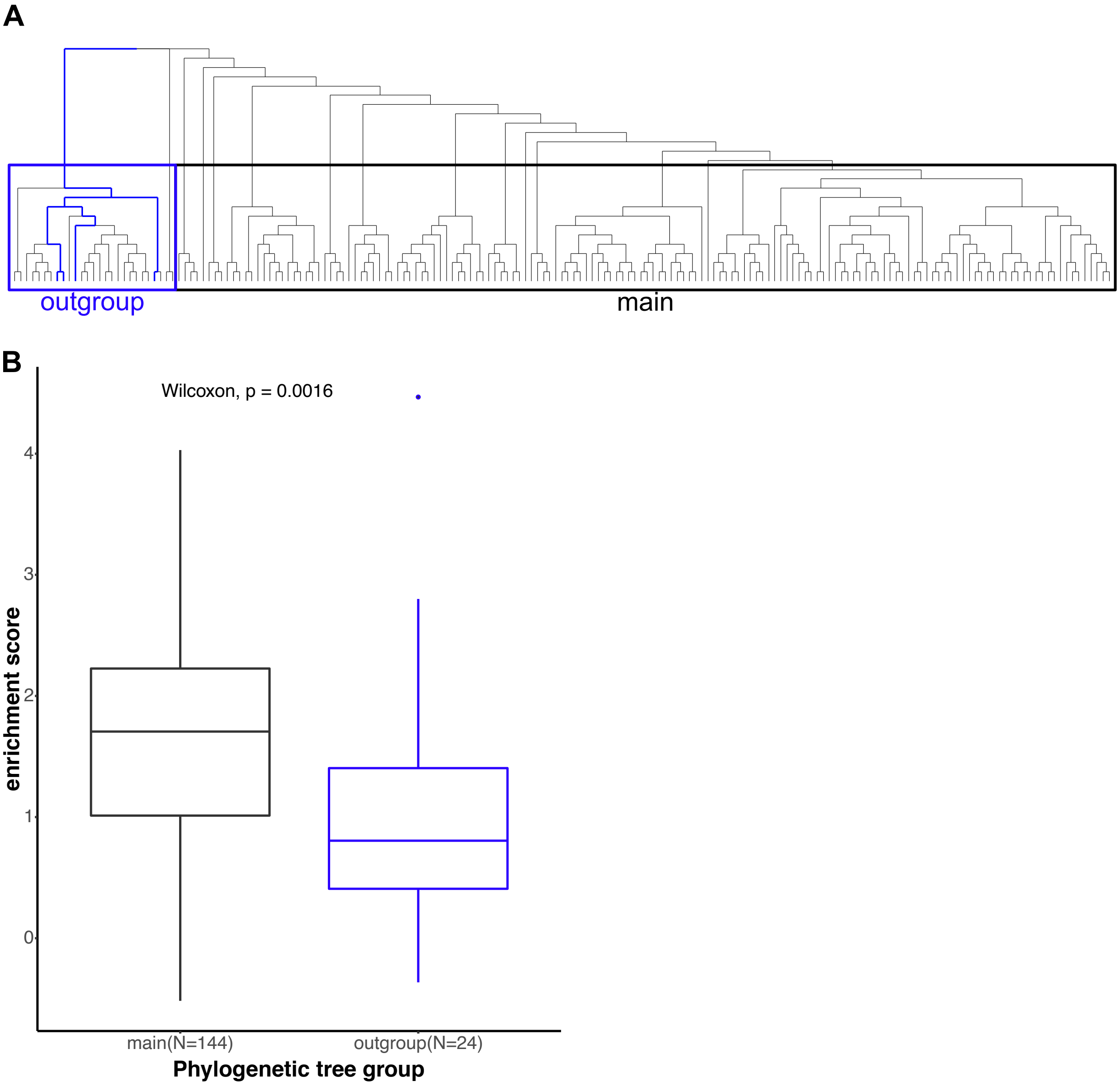


Supplemental Figure 7: Earliest diverging LTR18A lineage forms a functionally distinct outgroup. A) Phylogenetic tree of manual curated LTR18A splitting the earliest diverging lineage (outgroup, blue) from the main lineage (black). B) HepG2 enrichment scores of motif focused regions for main and outgroup hg19 LTR18A elements. *P* value derived from two-tailed Mann-Whitney *U* test.

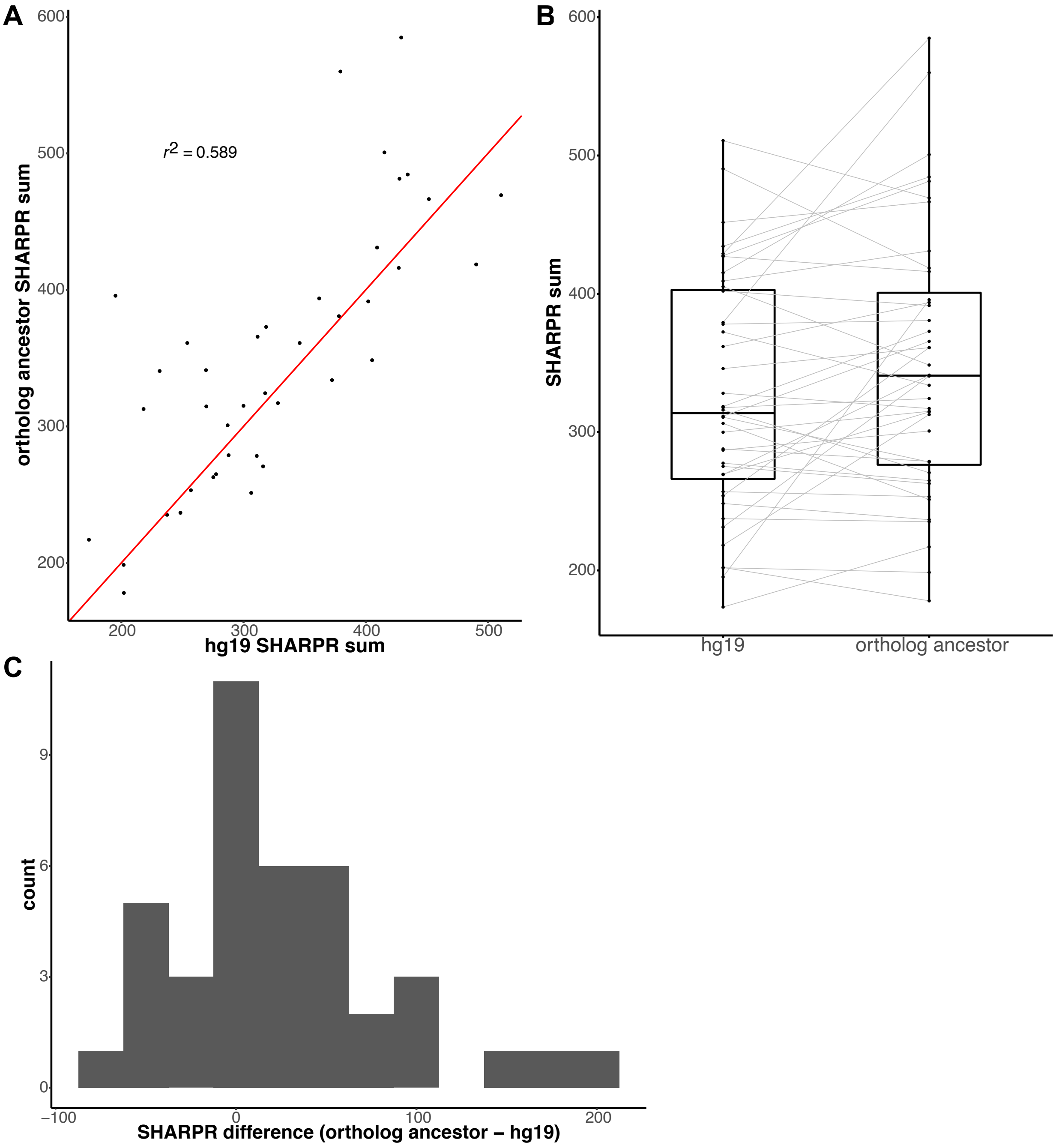


Supplemental Figure 8: Comparison of enhancer activities for present day hg19 LTR18A elements and their corresponding ortholog ancestors. A) Correlation of SHARPR sum for hg19 elements and their ortholog ancestors. The red line depicts a one-to-one relationship (y=x). B) Distribution of hg19 elements and ortholog ancestor SHARPR sums. Grey lines connect hg19 elements with their ortholog ancestor. C) Distribution of differences between ortholog ancestors and hg19 elements.

| | hg19 | panTro4 | gorGor3 | nomLeu3 | papAnu2 | rheMac3 | calJac3 |
|---|---|---|---|---|---|---|---|
| Expected (genome) | 0.00% | 1.37% | 1.74% | 3.76% | 6.16% | 6.15% | 10.08% |
| Expected (masked) | 0.00% | 1.45% | 1.86% | NA | 6.47% | 6.61% | 10.72% |
| Observed | 0.00% | 1.54% | 2.18% | 4.97% | 7.80% | 7.89% | 12.93% |
| Percent increase (observed over genome) | 0.00% | 12.61% | 25.17% | 32.39% | 26.76% | 28.13% | 28.18% |
| Number | 181 | 176 | 173 | 152 | 144 | 148 | 76 |

Supplemental Table 2: LTR18A substitution rate compared to genomic background. Expected substitution rates from genome-wide comparisons between hg19 and other primate genomes are shown for the entire genome and the masked (repetitive) genome. The observed rate is the average substitution rate for LTR18A orthologs between hg19 and other primate genomes. The number of orthologous LTR18A for each species is displayed on the last row.

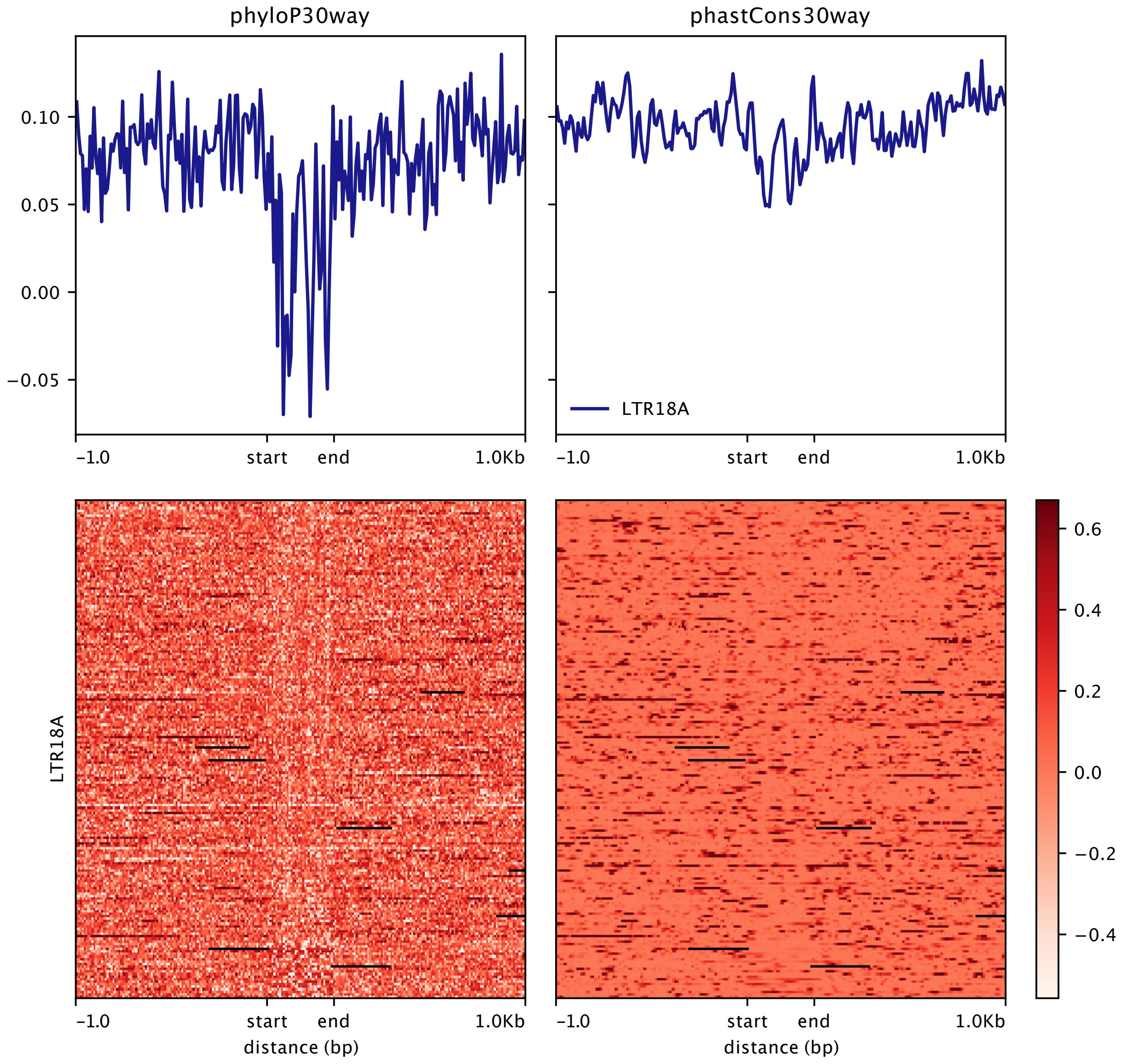


Supplemental Figure 9: Conservation scores for LTR18A and surrounding 1kb genomic regions. The phyloP and phastCons scores for 30 mammals (27 primates) are shown. Start and end refer to LTR18A start and end sites.

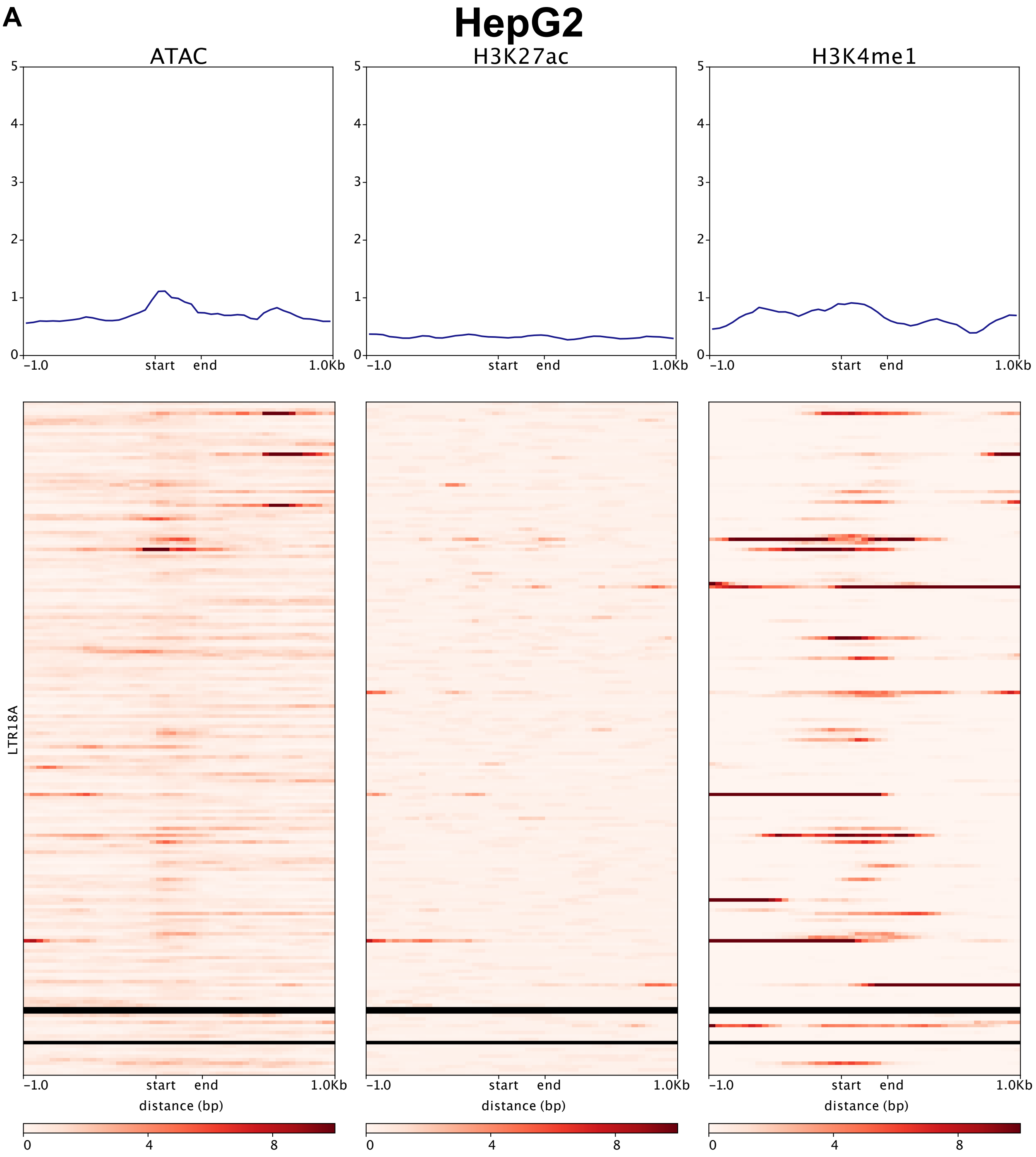
A
HepG2
ATAC
H3K27ac
H3K4me1
5
4
3
2
1
0
-1.0
start
end
1.0Kb
LTR18A
distance (bp)
0
4
8

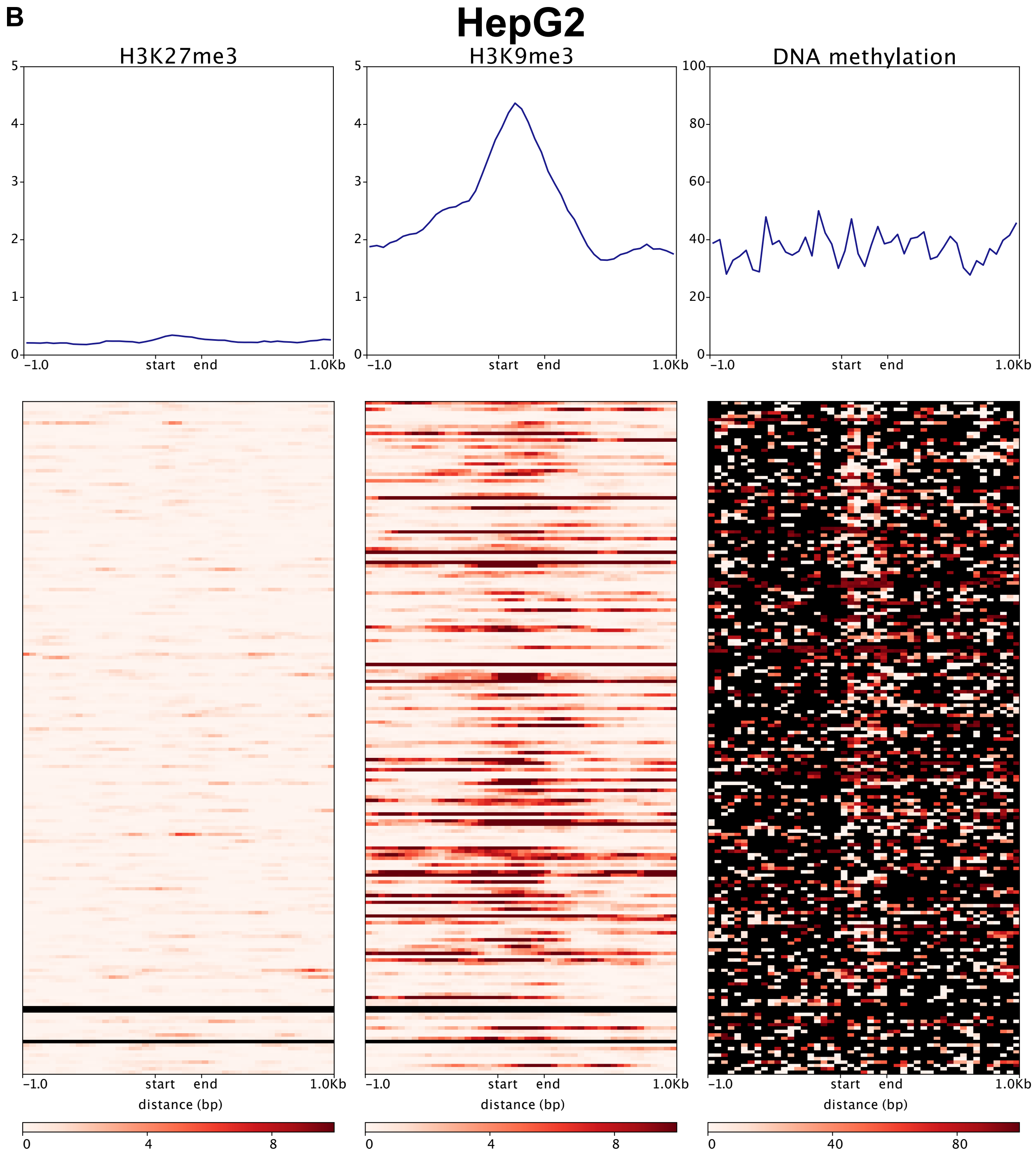
B
HepG2
H3K27me3
H3K9me3
DNA methylation
0
1
2
3
4
5
20
40
60
80
100
-1.0
start
end
1.0Kb
distance (bp)
0
4
8
0
40
80

C

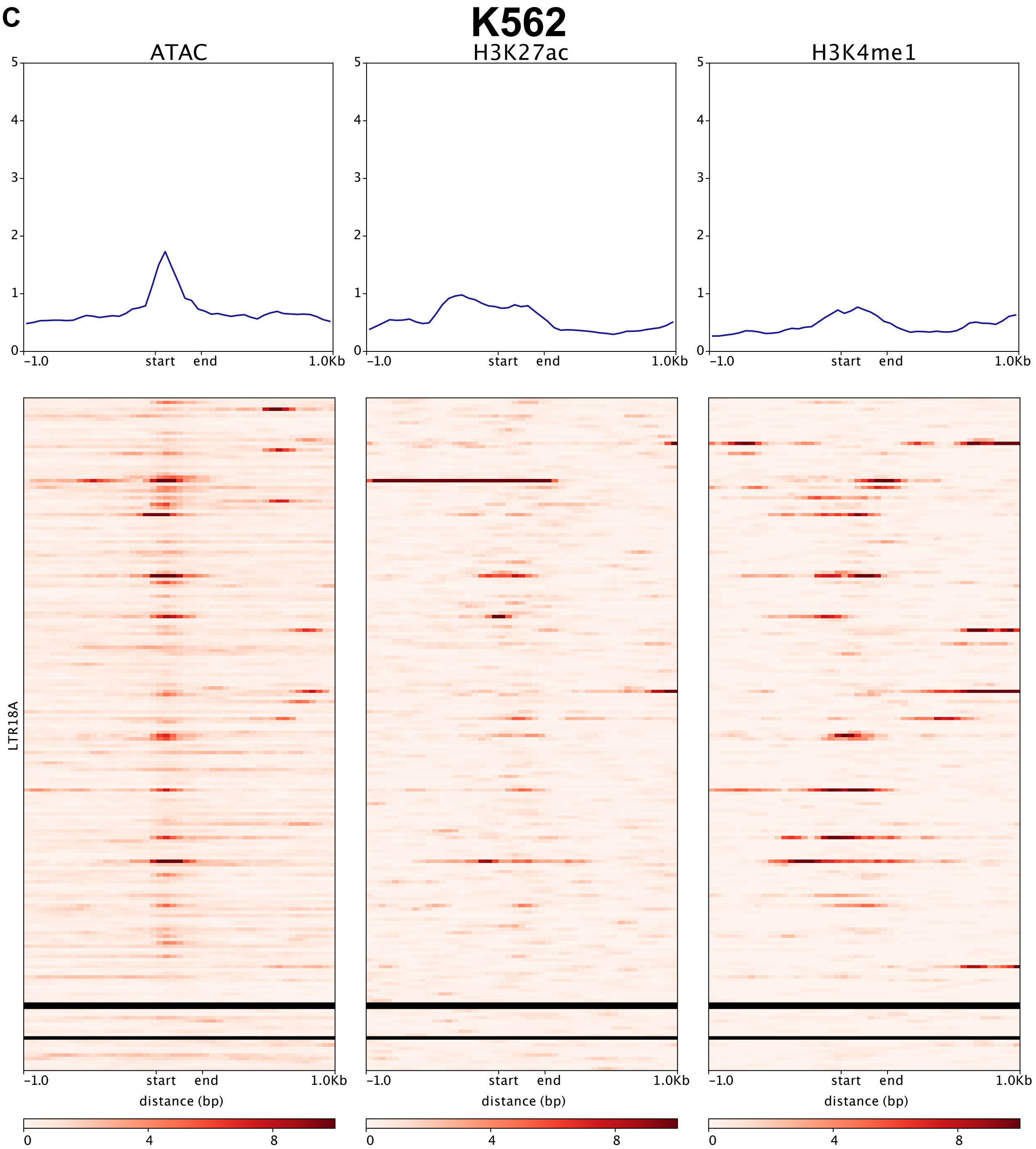

K562
ATAC
H3K27ac
H3K4me1
0
1
2
3
4
5
-1.0
start
end
1.0Kb
LTR18A
distance (bp)
0
4
8

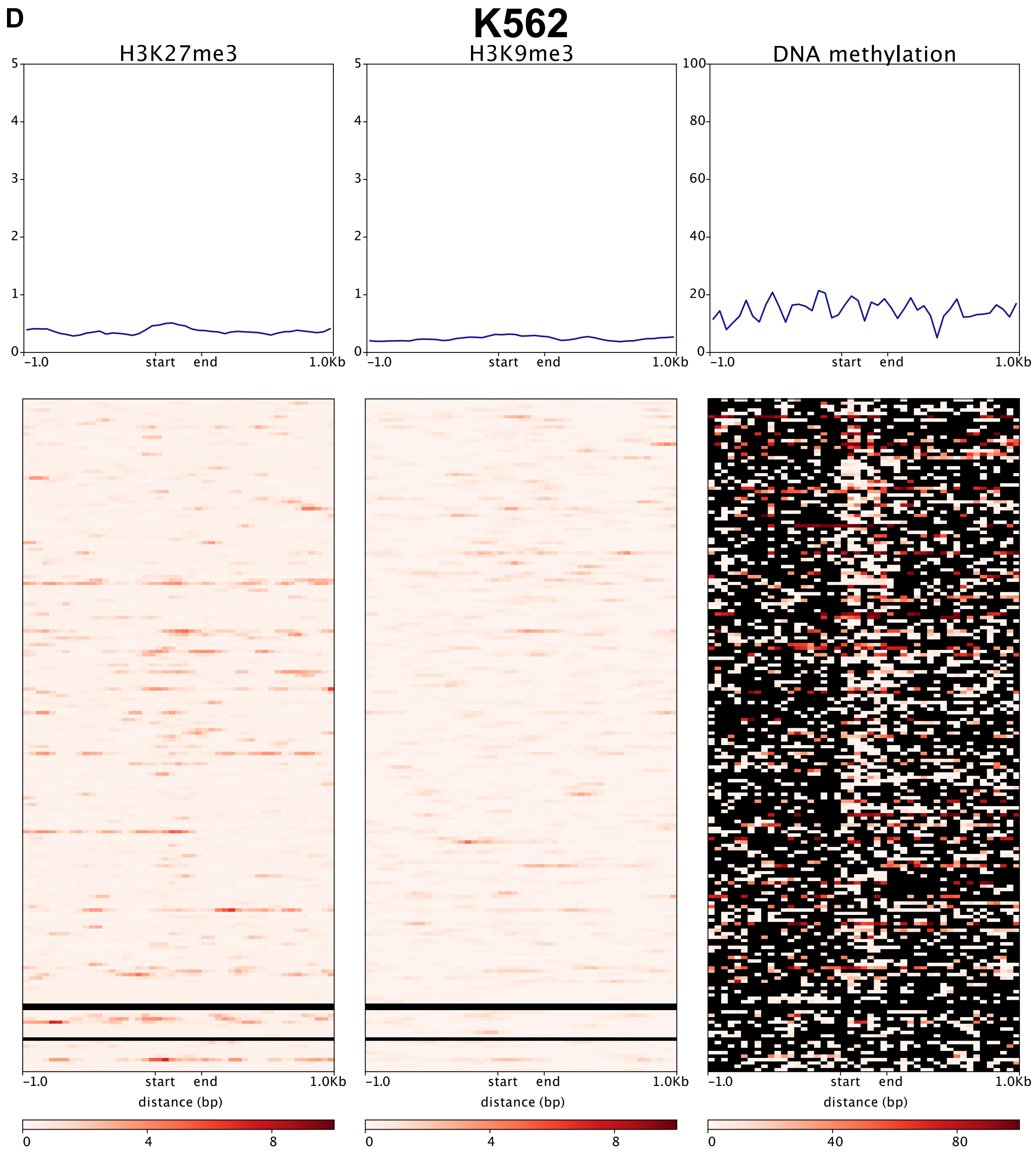


Supplemental Figure 10: Epigenetic marks at LTR18A in HepG2 (A and B) and K562 (C and D). Enhancer associated epigenetic marks (ATAC, H3K27ac, and H3K4me1) and repressive epigenetic marks (H3K27me3, H3K9me3, and DNA methylation) are shown. Three LTR18A elements have lengths shorter than the bin size (50bp) and are shown as black bars.
